# Spaceflight-relevant microgravity triggers a sublethal Parkinson’s-like dopaminergic decline in human neurons and organoids

**DOI:** 10.64898/2026.07.09.737494

**Authors:** Nilufar Ali, Shehbeel Arif, Max-Florian Mortimer, Matthew Brandner, Joshua Baldridge, Ruben Ferreira Paiva, Mukta S. Sane, Shin Pu, Luke Woodbury, Gunes Uzer, Cheryl Jorcyk, Greg Hampikian, Masihhur Laskar, Julia Oxford

## Abstract

Spaceflight stressors may increase Parkinson’s disease (PD) risk, but microgravity’s specific contribution to human dopaminergic (DA) vulnerability remains undefined. Here, we exposed human iPSC-derived midbrain DA organoids and differentiated SH-SY5Y neurons to simulated microgravity for 72 hours. This exposure reduced neurite outgrowth, eroded DA identity, and activated familial-PD mitochondrial kinases without depleting extracellular dopamine. We observed severe mitochondrial dysfunction—including membrane potential loss and respiratory suppression—coupled with global translational repression. Furthermore, a sublethal, pre-degenerative state emerged, characterized by the selective release of mitochondrial cell-free DNA without apoptotic activation. Proteomic profiling revealed striking convergence with human PD transcriptomes and astronaut blood, highlighting shared suppression of mitochondrial metabolism, ribosomal translation, and altered RNA splicing. Together, these findings establish simulated microgravity as a sufficient, non-toxin trigger of early PD-like DA dysfunction, providing a robust human model for investigating prodromal neurodegeneration.

## Introduction

Parkinson’s disease (PD) was clinically described more than two centuries ago, yet the molecular events that initiate DA neuronal degeneration remain an active area of research. With the global aging population and PD incidence rising, identifying tractable mechanisms and modifiable risk factors has become increasingly urgent^1,2^. Fewer than 10% of cases can be ascribed to monogenic causes, leaving the vast majority, termed idiopathic PD, thought to arise from complex interactions involving aging, genetic susceptibility and life-long exposures to agents including pesticides, heavy metals and ionizing radiation^3^. Aging remains the strongest risk factor, characterized by molecular hallmarks including oxidative stress, mitochondrial dysfunction and impaired proteostasis, that converge with nigrostriatal degeneration^4–6^.

Spaceflight imposes a constellation of extreme environmental stressors. These include microgravity, ionizing cosmic radiation, hypercapnia, sleep disruption, and social isolation, features in common with accelerated aging that have been proposed as drivers of PD-like changes in DA neurons during missions exceeding ∼35 days^7,8^. Candidate mechanisms include disrupted mitochondrial bioenergetics, elevated reactive oxygen species (ROS), impaired autophagy-lysosome function and dysregulation of the integrated stress response (ISR)^7^. Consistent with this, stimulated microgravity (SMG) alters brain structure and oxidative balance in rodent models in a region- and duration-dependent manner^9^. However, the contribution of microgravity itself, uncoupled from radiation and other co-stressors, to DA vulnerability in human cells has not been systematically studied.

A major limitation in the field is that mechanistic insight into spaceflight-induced neurological change has so far depended on peripheral biospecimens (blood, fibroblasts, and hair follicles) from astronauts or post-mortem tissue from rodent missions aboard the International Space Station (ISS). This impairs causal interrogation of the human DA lineage and precludes scalable therapeutic screening^7^. Human induced pluripotent stem cell (iPSC)-derived neural organoids offer a tractable alternative, recapitulating human midbrain cytoarchitecture and supporting neuronal maturation in three dimensions^10–13^. Such organoids have recently been deployed in low-Earth orbit and under simulated microgravity, demonstrating their suitability for studying nervous-system responses to spaceflight stressors^14,15^.

In this study, we isolated the contribution of microgravity to DA vulnerability using a ground-based simulation. Human iPSC-derived midbrain DA organoids and adherent 2D differentiated SH-SY5Y DA neurons were exposed to SMG on a tunable custom-built clinostat. We show that 72 h of SMG is sufficient to induce an early PD-like molecular signature in the DA system across both 2D and 3D models. These findings establish ground-based SMG exposure of human DA cultures as a physiologically relevant, toxin-free platform for modeling microgravity induced changes in space and interrogating early PD-like molecular stressor and for screening neuroprotective interventions.

## Results

### Clinostat optimization to achieve microgravity

To quantify the simulated microgravity conditions produced by the clinostat (Fig 1A, B), acceleration data were collected from both the innermost and outermost positions at multiple rotation speeds (Fig 1C, D). Using a tri-axial accelerometer outputs in Euclidean norm^16^ (Fig 1E) we show the mean acceleration vector components and the corresponding net acceleration (a) at the center-most position (Fig 1F, G). At 15 RPM (the rotation rate at which all adherent cell experiments were conducted)^17^ the clinostat produced an effective time-averaged randomization of the gravity vector, with a residual acceleration of approximately 0.048 g (∼0.47 m s⁻²) at the center position. Although this does not constitute true microgravity, these operating conditions are widely used to simulate aspects of the reduced gravitational loading experienced during spaceflight by minimizing the net directional influence of gravity on cultured cells. Increasing the rotation speed to 25 RPM at this position led to a slight rise in experienced acceleration to 0.086 g, suggesting a decrease in the quality of vector canceling due to centrifugal effects.

**Figure 1:**
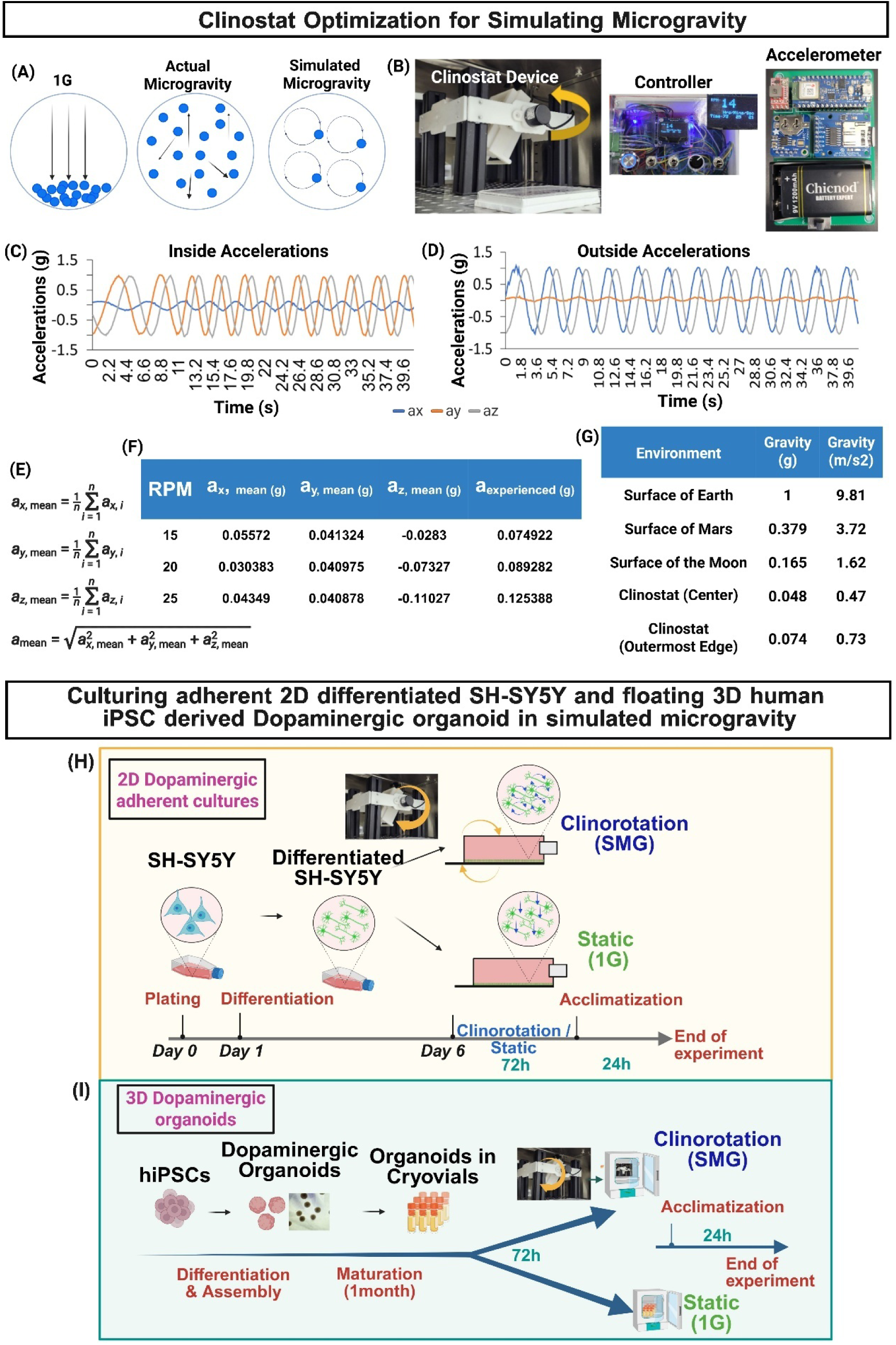
Clinostat Optimization. (A) shows principle of clinorotation induced microgravity (B) Operational components of our clinostat. (C) Sinewave of the inside collected acceleration for the first 30 second interval depict the dynamic acceleration waveforms over a 30-second interval for (D) Sinewave of the outside collected acceleration for the first 30 second interval. The period and amplitude of the sine wave oscillations confirm the expected rotation dynamics of the clinostat. Acceleration table of acceleration vectors collected by the Aurdiuno accelerometer on the innermost edge of the clinostat. (E) Equation used to calculate mean acceleration (F) Optimization of different RPMs leading to experienced accelerations. (G) Gravity as experienced on Earth, Mars, and the Moon compared to positions on the clinostat at 15 RPM. Paradigm of culturing 2D adherent differentiated SH-SY5Y DA neurons (H) and one month old human iPSC derived DA brain organoids (I) in optimized microgravity simulator for 72h.

In contrast, (Fig 1G). shows the acceleration data collected from the outermost edge of the clinostat. Here, the experienced gravity was consistently higher than at the center, with values ranging from 0.075 g at 15 RPM to 0.125 g at 25 RPM. This increase is attributable to the larger radial distance from the axis of rotation, which increases centrifugal forces. Despite this, the values still remain well below Earth’s gravity (1 g), indicating that the outer edge can still approximate partial gravity environments, such as those found on the Moon (0.165 g) or Mars (0.379 g), as summarized in (Fig 1G).

For experiments, rotation speeds were selected to balance effective gravity vector randomization while minimizing centrifugal acceleration and shear stress. Previous clinostat studies have demonstrated that simulated microgravity is achieved through continuous reorientation of the gravity vector, resulting in time-averaged nullification of gravitational loading on cells^18,19^.

Previous work adapting 2D clinostats for adherent cells demonstrated that rotational speed must be optimized to reduce residual g-forces while maintaining appropriate culture stability and minimizing shear-induced effects^20,21^. For adherent cell cultures, 15 RPM was selected to ensure sufficiently rapid reorientation of the gravity vector across the attached monolayer while avoiding excessive fluid motion and centrifugal artifacts^17^. For organoid cultures, lower rotational speeds (10-12 RPM) were used to minimize residual acceleration and fluid shear while maintaining adequate suspension of the organoids within the culture medium^22,23^. This range is consistent with rotational conditions commonly used for low-shear organoid and rotating culture systems, where approximately 10–15 RPM has been shown to support tissue integrity and nutrient exchange while reducing mechanical stress. Since centrifugal acceleration increases proportionally with the square of angular velocity: a_c_=rω^2^ higher rotational speeds can introduce increasing centrifugal and shear forces that may confound biological responses. Therefore, RPMs were intentionally kept at the lowest values that maintained effective gravity vector averaging and sample stability throughout the experiments.

### Simulated microgravity reduces neurite length and organoid sprouting mimicking an early hallmark of neurodegeneration

To assess the impact of simulated microgravity (SMG) on neuronal morphology, differentiated SH-SY5Y neurons were exposed to SMG or 1G for 72 hours. Compared to 1 G controls, SMG-treated neurons exhibited a reduced neuronal population (Fig. 2A). Quantitative analysis revealed a significant decrease in neurite length under SMG (Fig. 2B, C), whereas neurite width remained unchanged (Fig. 2B, D), indicating selective impairment of neurite extension rather than gross cytoskeletal alteration. Evaluation of cytotoxicity via LDH assay indicated increased LDH activity in SMG group compared to control resulting in a mild ∼9.8% neuronal toxicity (Fig 2E, F).

**Figure 2:**
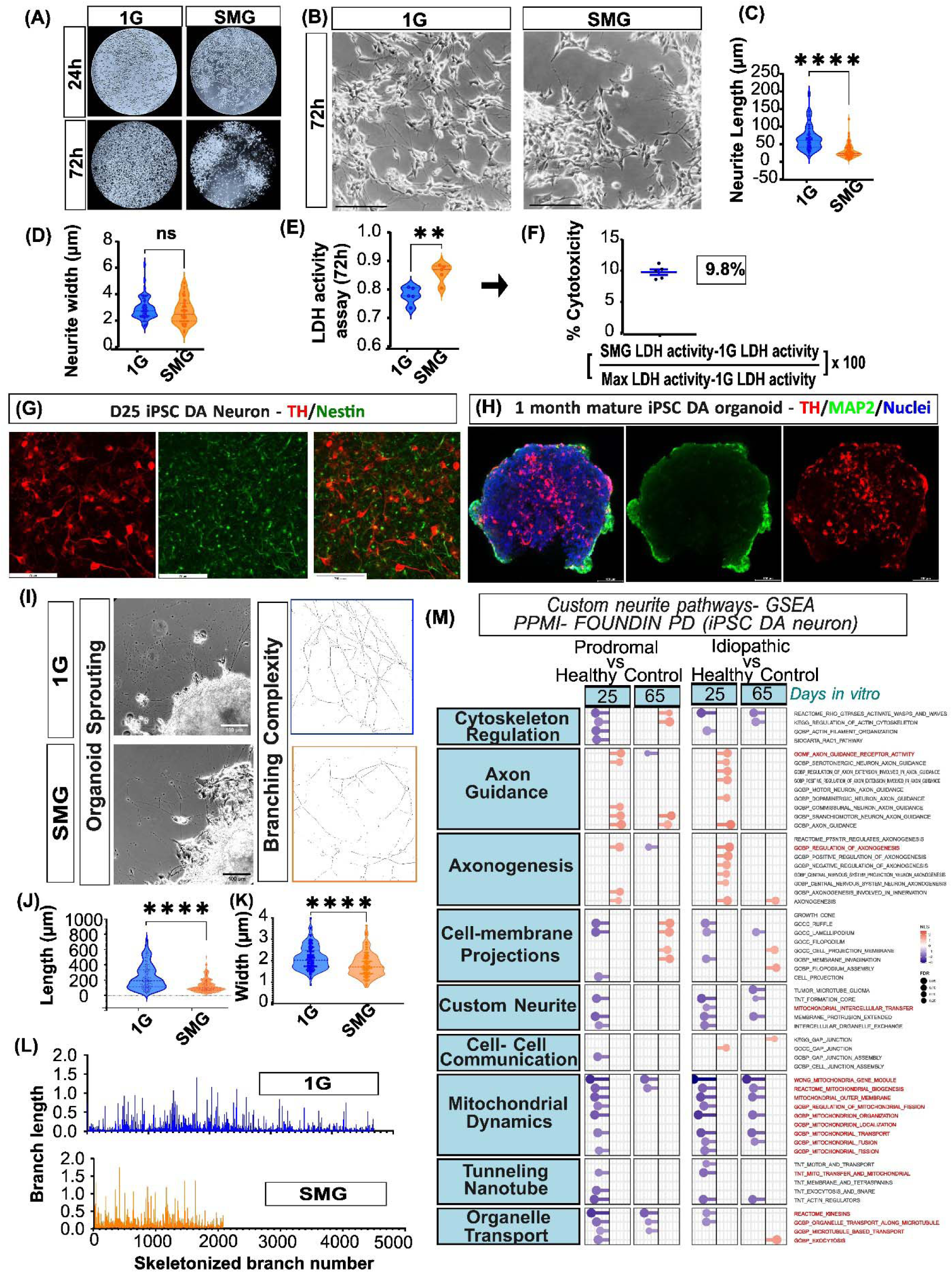
Simulated microgravity impairs neurite outgrowth and organoid sprouting. (A) Representative phase-contrast images of retinoic acid–differentiated SH-SY5Y neurons under control (1 G) and simulated microgravity (SMG) conditions at 24h and 72 h, showing reduced neuronal population under SMG. (B) Representative images showing neurite extensions from neurons cultured at 1G and in SMG for 72h. Scale bar: 20 µm. (C) Quantification of neurite length (µm) showing a significant reduction under SMG and (D) neurite width (µm), showing no significant difference between control and SMG groups (p < 0.05, unpaired t-test). Data represent mean ± SEM from n = 6-14. (E, F) LDH assay quantification among groups indicating ∼9.8% cytotoxicity in neurons post-SMG, n = 5. (G) Immunofluorescence of human iPSC–derived DA organoids stained for tyrosine hydroxylase (TH), Nestin positive neuron at 25 div (day 1 - before assembling organoid) and (H) TH/Map2 positive at 30 days mature organoid, demonstrating robust DA neurons. Scale bar: 100 µm (I) Neurite sprouting from 1G and SMG exposed organoids, showing neurite network and branching complexity. Scale bar: 100 µm. (J) Quantification of neurites show a significant reduction in their length (µm) under SMG as well as (K) width, when compared to control groups. n = 6-14. (L) Analysis of branching complexity reveal reduction in neurite branching and length in SMG group as compared to 1G. (M) GeneSet enrichment analysis (GSEA) using custom curated neurite pathways consisting cytoskeletal regulation, cell-membrane projection, axon guidance, axonogenesis, cell-cell communication, mitochondrial dynamics, tunneling nanotube and organelle transport pathways on PD patients vs control iPSC derived DA neruons at DIV 25 and DIV 65 obtained from MJFF PPMI Foundin PD bulk RNA seq dataset. GSEA performed on Prodromal vs healthy control and idiopathic PD vs healthy control. False discovery rate (FDR) shown in lolipop sizes ranging from 0.2 to 0.05 and normalized enrichment score for pathways shown as heatmap ranging from −3 (blue) to +2 (red) depicting downregulated and upregulated pathways respectively. All experimental data represent mean ± SEM followed by two-tailed unpaired Student’s t-test, *p < 0.05 considered statistically significant.

While 2D monocultures provide a foundational baseline for neuronal stress, they lack the complex cytoarchitecture and spatial interactions of the human midbrain. To determine if these early morphological deficits are conserved in a three-dimensional tissue context, we generated human iPSC-derived DA organoids expressing Nestin/TH-positive neurons (Fig. 2G). When subjected to SMG for 72h, the organoids showed a reduction in size (Fig. 2H). Post SMG or 1G exposure these organoids were assessed for neurite sprouting which revealed that control 1 G organoids generated extensive neurite networks, whereas SMG-exposed organoids displayed markedly reduced sprouting and branching complexity (Fig. 2I). Quantification of the images revealed a significant decrease in both length and width of the projections (Fig 2J, K) as well as a decline in branching complexity measured by reduced branches and their length in SMG group (Fig2 L). To validate our neuronal projection, decline as early PD event, we performed geneset enrichment analysis (GSEA) using custom curated neurite pathways consisting cytoskeletal regulation, cell-membrane projection, axon guidance, axonogenesis, cell-cell communication, mitochondrial dynamics, tunneling nanotube and organelle transport pathways on MJFF PPMI Foundin PD dataset. We compared Prodromal vs healthy control and idiopathic PD vs healthy control at day 25 and day 65 of differentiation (Fig 2M). Our analysis revealed a highly significant (FDR 0.05) downregulation of cytoskeletal regulation pathways as an effect of Prodromal PD as well as established idiopathic PD when compared to controls. Additionally, cell membrane projections, neurite, tunneling nanotubes related pathways were also inhibited in Prodromal iPSC derived DA neuron and they became increasingly enriched in established Idopathic PD patients vs control at both days in vitro. Axonogeneis pathways however were upregulated in both prodromal and idopathic Pd groups revealing compensatory signaling. Thus, the findings elude to fact that microgravity induces a mild and early PD like pathology.

### Simulated Microgravity upregulates familial Parkinson’s Disease–Associated Stress Pathways

In 2D, adherent DA neuronal system-differentiated SH-SY5Y, simulated microgravity (SMG) resulted in a significant loss of DA molecular identity accompanied by activation of Parkinson’s disease–associated stress pathways. Quantitative RT-PCR analysis demonstrated a downregulation of Tyrosine Hydroxylase (TH) and alpha synuclein mRNA, alongside increased expression of PTEN-induced putative kinase 1 (PINK1), Parkin RBR E3 ubiquitin protein ligase (PRKN), and Leucine-rich repeat kinase 2 (LRRK2) following SMG exposure (Fig. 3A–E). This transcriptional pattern mirrors alterations previously reported in astronaut peripheral blood^7^, suggesting conserved microgravity-responsive neurodegenerative signaling. Consistent with transcriptomic findings, Western blot analysis revealed a reduction in TH protein levels under SMG conditions, which was confirmed by densitometric quantification (Fig. 3F, G). In contrast, dopamine neurotransmitter levels remained unchanged between groups, suggesting the presence of compensatory mechanisms in surviving DA neurons to maintain dopamine homeostasis despite reduced TH expression. Such compensatory preservation of dopamine signaling has previously been reported during early DA dysfunction and neurodegeneration^24,25^. Alternatively, SMG conditions may enhance dopamine release or alter vesicular exocytosis dynamics, thereby maintaining extracellular dopamine levels despite reduced biosynthetic capacity.

**Figure 3.**
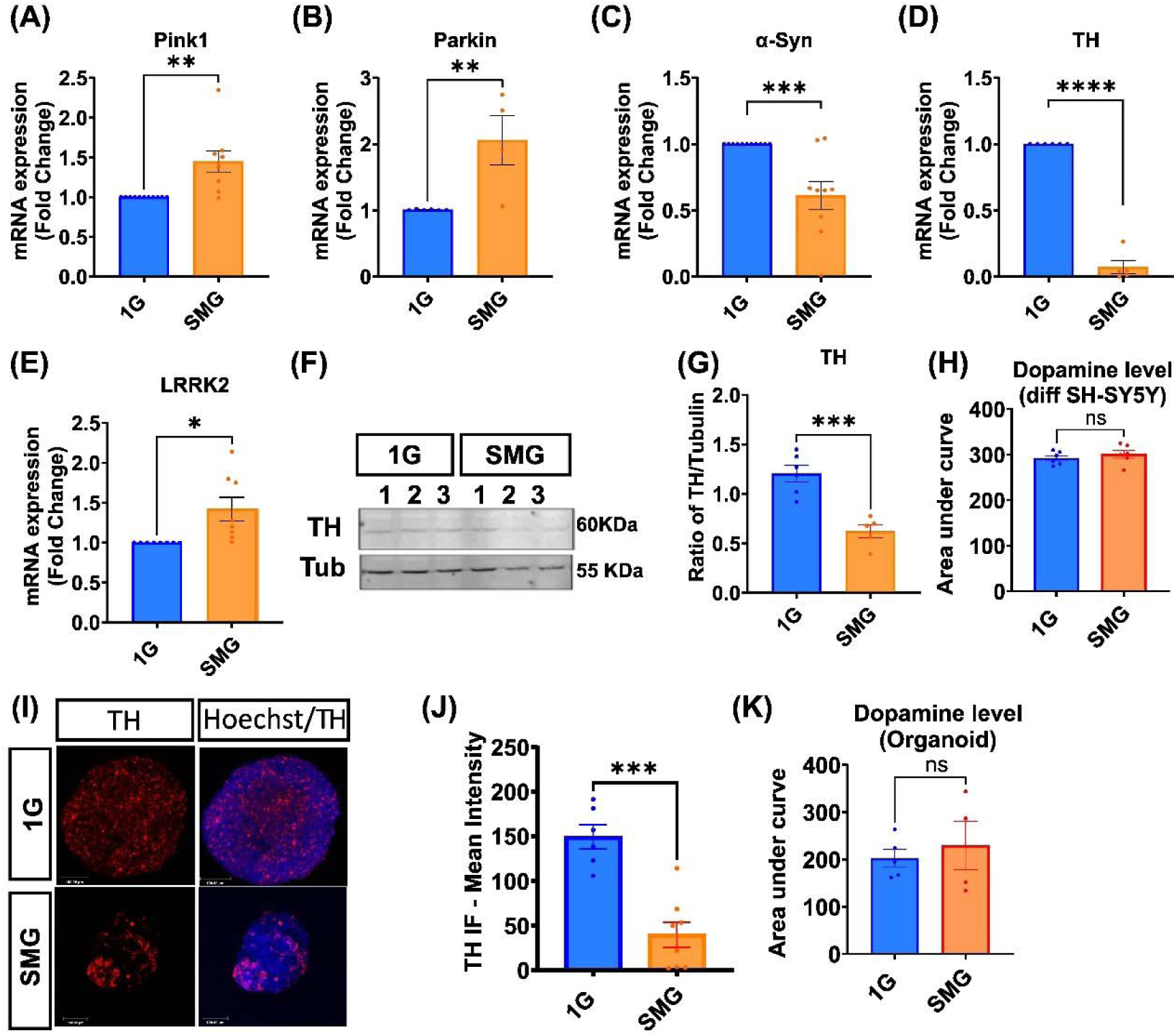
Simulated microgravity induces DA molecular and functional deficits in human neuronal models. (A–E) Quantitative RT-PCR analysis of differentiated SH-SY5Y neurons showing reduced Tyrosine Hydroxylase (TH) mRNA and increased expression of PTEN-induced putative kinase 1 (PINK1), Parkin RBR E3 ubiquitin protein ligase (PRKN), and Leucine-rich repeat kinase 2 (LRRK2) following simulated microgravity (SMG) exposure relative to 1G controls. n=3-8. (F) Representative Western blot images showing decreased TH protein expression under SMG conditions. (G) Densitometric quantification of TH protein expression normalized to loading control Tubulin (Tub) n= 4-8. (H) Mass spectrometry-based quantification of spontaneous dopamine neurotransmitter release from culture medium post 1G or SMG exposure. n=5-6. (I) Immunofluorescence and (J) mean intensity quantification of human iPSC– derived DA organoids stained for tyrosine hydroxylase (TH, red) and nuclei (Dapi, blue) post 1G or SMG exposure (n=6-8). (K) Mass spectrometry-based quantification of spontaneous dopamine neurotransmitter release from culture medium of brain organoids post 1G or SMG exposure. n=5. All experimental data represent mean ± SEM followed by two-tailed unpaired Student’s t-test, *p < 0.05 considered statistically significant.

### Human DA neurons under simulated microgravity exhibit mitochondrial dysfunction

Based on our previous findings that spaceflight stressors induce mitochondrial dysfunction^7^, we investigated the effect of simulated microgravity (SMG) on mitochondrial function in DA neurons. DA organoids exhibited a significant decrease in mitochondrial membrane potential, measured by TMRM staining, following 30 h of SMG exposure (Fig. 4A, C).

**Figure 4.**
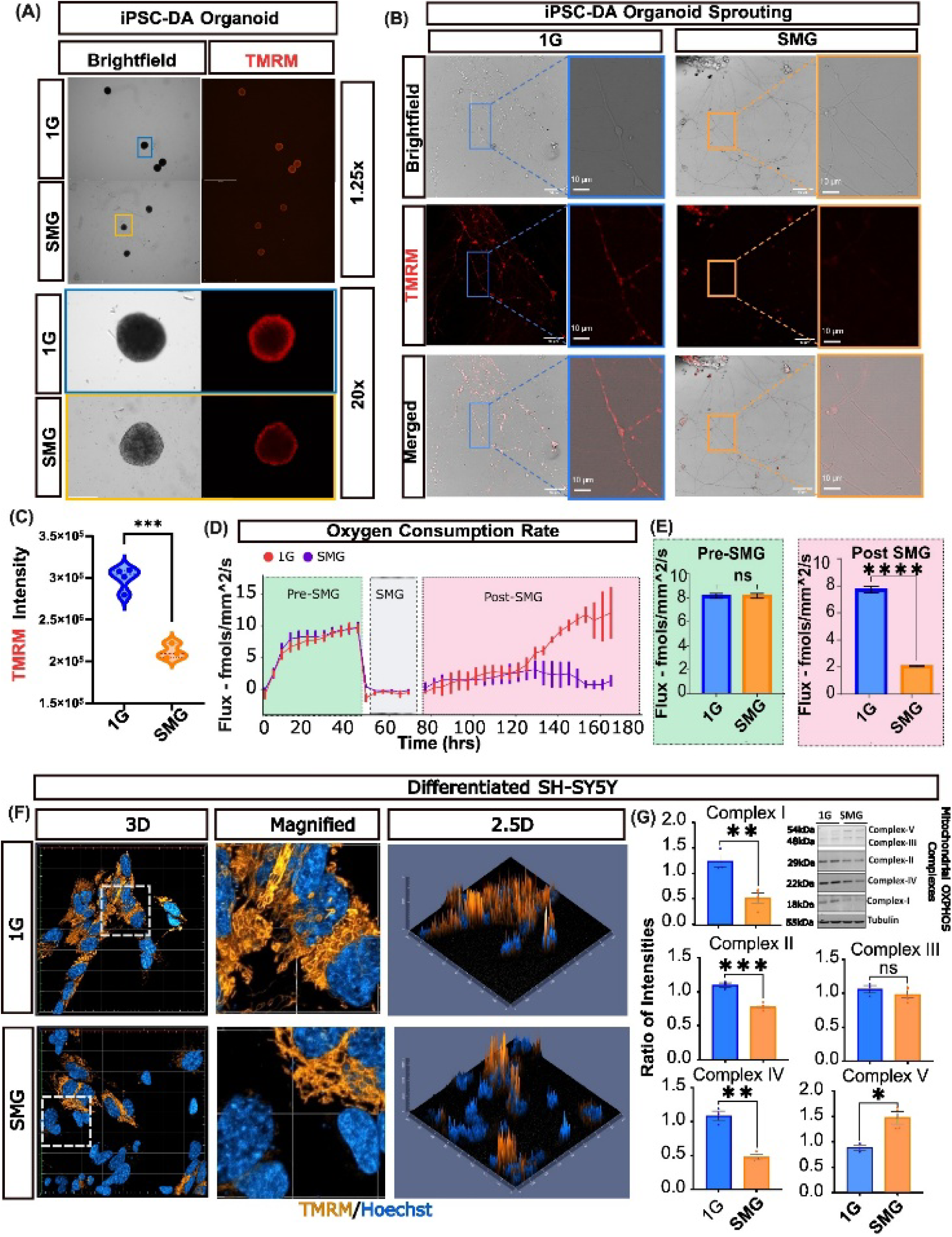
Simulated microgravity induces mitochondrial dysfunction in neuronal models. (A) Fluorescence microscopy images (1.25× and 20× magnification) of human iPSC-derived DA organoids following simulated microgravity (SMG) exposure. SMG organoids exhibit reduced mitochondrial membrane potential, measured by tetramethylrhodamine methyl ester (TMRM) staining, compared to 1G controls. (B) Confocal images of sprouted organoids under 1G and SMG conditions showing neurite projections. In 1G, neurites display robust TMRM-positive mitochondria, whereas SMG-exposed organoids show a marked reduction in mitochondrial signal within projections. Scale bar: 10 µm. (C) Quantification of TMRM fluorescence intensity demonstrating a significant decrease in mitochondrial membrane potential in SMG compared to 1G controls (N = 3-4). (D) Continuous measurement of organoid respiration assessed by oxygen consumption rate (OCR; fmol/mm²/s). Organoids displayed comparable baseline respiration prior to SMG exposure. Following ∼24 h of SMG, a pronounced decline in OCR was observed in the SMG group, whereas 1G controls recovered and maintained normal respiratory activity (N = 4) (E) Quantification of OCR showing no significant difference between groups at baseline (pre-SMG), while post-treatment SMG exposure resulted in a significant reduction in OCR, indicating metabolic suppression. (F) Differentiated SH-SY5Y neuronal cultures exposed to SMG exhibit reduced mitochondrial membrane potential compared to 1G controls, as assessed by TMRM fluorescence. Representative fluorescence images and 2.5D projections reveal uniform mitochondrial distribution associated with nuclear signal in 1G conditions, whereas SMG conditions show heterogeneous mitochondrial signal intensity, reduced mitochondrial membrane potential, and occasional punctate high-intensity TMRM signals, indicative of mitochondrial heterogeneity and dysfunction. (G) Western blot analysis of mitochondrial oxidative phosphorylation (OXPHOS) complexes in differentiated SH-SY5Y neurons. SMG exposure results in significant reductions in Complex I, II, and IV protein levels, with no significant change in Complex III. Complex V (ATP synthase) is upregulated, suggesting a potential compensatory response to impaired electron transport chain function. (N = 3-5, data presented as mean ± SEM). All experimental data represent mean ± SEM followed by two-tailed unpaired Student’s t-test, *p < 0.05 considered statistically significant.

In addition, sprouted SMG-exposed organoids showed a marked reduction in TMRM-positive mitochondria within neuronal projections compared to 1G controls (Fig. 4B), indicating impaired mitochondrial presence and/or transport along axonal processes. This observation is consistent with previous MJFF FOUNDIN-PD iPSC-derived DA neuron analyses highlighting mitochondrial dynamics and organelle transport pathway disruptions (Fig. 2M).

We next assessed real-time continuous organoid respiration before and after SMG exposure (Fig. 4D). Following an initial growth phase (0–45 h), organoids were removed from the plate at ∼45 h and subjected to their respective gravity conditions ex situ for ∼30 h. Organoids were then re-seeded at ∼75 h for continued monitoring, and the experiment was conducted over 7 days (168 h total). Under baseline conditions, 1G and SMG organoids exhibited comparable respiratory activity, reaching ∼10 fmol/mm²/s with overlapping SEM intervals during the early growth phase. However, post-recovery dynamics diverged significantly. 1G organoids resumed metabolic activity more rapidly, reaching positive flux earlier and continuing to increase to ∼12.3 fmol/mm²/s by 168 h. In contrast, SMG organoids exhibited delayed recovery and plateaued at ∼0.9 fmol/mm²/s, representing an approximately 92% sustained reduction in metabolic output relative to 1G controls.

To exclude the possibility that observed effects were due to mechanical stress or shear forces, we examined adherent 2D cultures of differentiated SH-SY5Y DA neurons exposed to SMG or 1G conditions. Consistent with organoid data, SMG exposure in these neurons resulted in a significant reduction in TMRM signal compared to controls.

Further analysis revealed increased heterogeneity in mitochondrial membrane potential within the SMG group. High-resolution 2.5D projections demonstrated an increased proportion of nuclei associated with diminished mitochondrial signal intensity, indicating impaired mitochondrial function at the single-cell level following SMG exposure.

To further characterize mitochondrial integrity, we assessed oxidative phosphorylation (OXPHOS) complexes by immunoblotting. Western blot analysis revealed significant reductions in mitochondrial Complex I, II, and IV protein levels in SMG-exposed cells, while Complex III remained largely unchanged. Notably, Complex V (ATP synthase) was upregulated, potentially reflecting a compensatory response to impaired electron transport chain activity (Fig. 4B). These changes corroborate with PD models in human DA neurons and in PD patients^26,27^.

Collectively, these findings demonstrate that simulated microgravity induces robust mitochondrial dysfunction across both neuronal cell lines and human iPSC-derived DA organoids, suggesting a conserved vulnerability of neuronal mitochondrial networks to altered gravitational conditions.

### Spaceflight and PD datasets reveal convergent translational suppression in DA neurons

Microgravity not only induce mitochondrial dysfunction but also alter ribosomal pathways and translation, as previously reported in human cell lines, Caenorhabditis elegans, and Arabidopsis thaliana^28–30^. Previously, we reported that oxidative phosphorylation (OXPHOS)–related genes— pathways also disrupted in PD—are significantly altered in astronaut blood following spaceflight^7^.

To further investigate this, we analyzed blood transcriptomics datasets from the Parkinson’s Progression Markers Initiative (PPMI) and the Parkinson’s Disease Biomarkers Program (PDBP), from previously published data^31^. We reanalyzed these datasets by comparing PD patients with healthy controls and performed gene set enrichment analysis (GSEA) using KEGG and MSigDB Hallmark pathways (Fig 5 A-D). Across both datasets, ribosomal pathways, ribosome biogenesis, and mitochondrial pathways emerged as some of the most significantly altered pathways.

**Figure 5:**
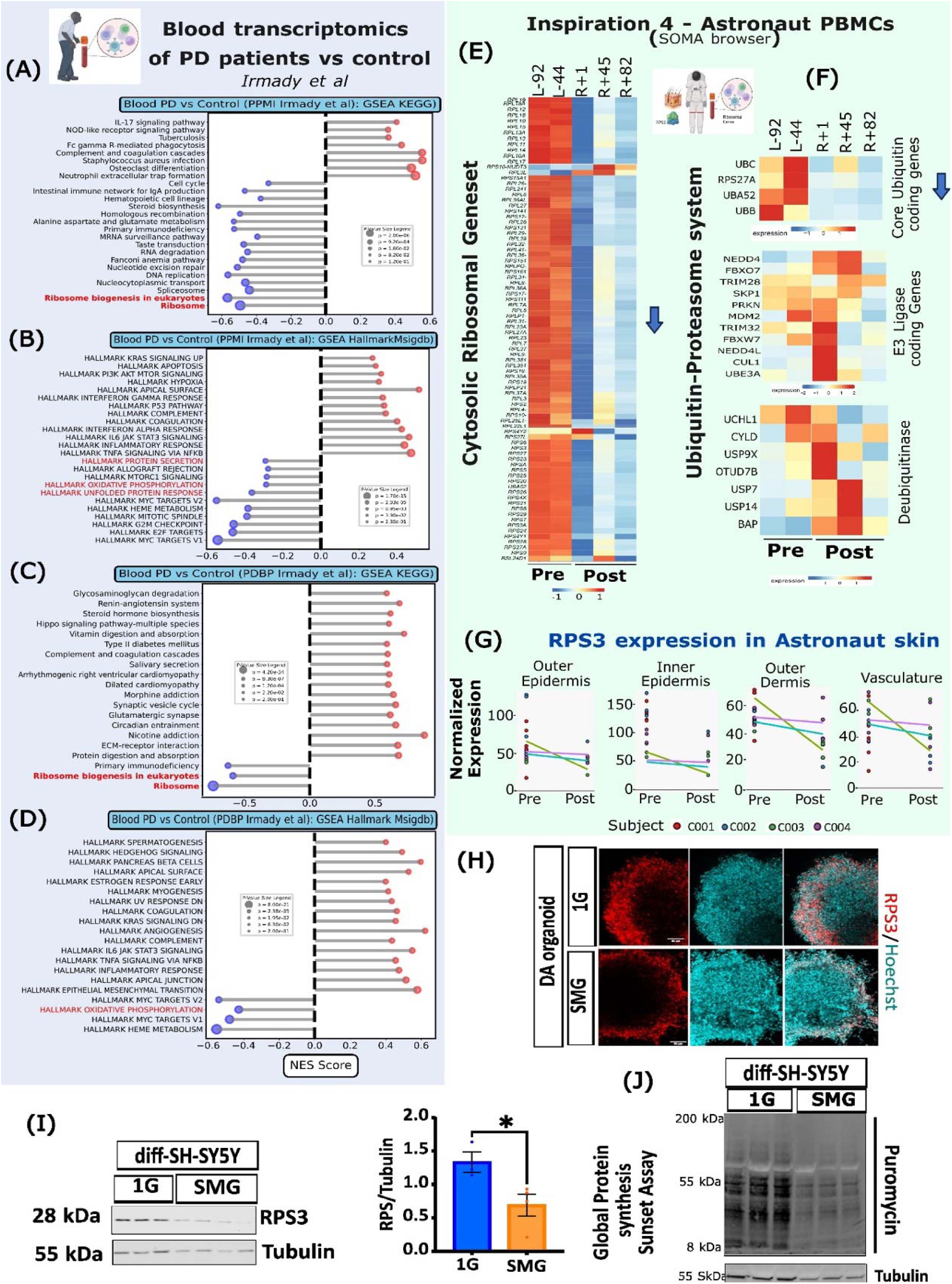
Convergent loss of Ribosomal pathways in PD blood and Astronaut blood PBMC post Spaceflight. (A–D) Gene set enrichment analysis (GSEA) of blood transcriptomic datasets comparing PD patients and healthy controls. Data were reanalyzed from Irmady et al^31^ using the Parkinson’s Progression Markers Initiative (PPMI-A, B) and Parkinson’s Disease Biomarkers Program (PDBP-C, D) cohorts. Lolipop plots showing normalized enrichment score (NES) on X axis and p value as lolipop sizes for GSEA using KEGG and MSigDB Hallmark pathways. Significant downregulation observed across both datasets-PPMI and PDBP cohorts in oxidative phosphorylation, ribosomal pathways, ribosome biogenesis, protein secretion, spliceosomal pathways, and related mitochondrial processes. (E–G) Transcriptomic analysis of astronaut peripheral blood mononuclear cells (PBMCs) comparing pre-flight and post-flight conditions. (E) Data (Soma browser analysis)^32^ show a sustained downregulation of cytosolic ribosomal gene sets following spaceflight, observed from return day +1 through up to 82 days post-flight. (F) Analysis of the ubiquitin–proteasome system in astronaut blood revealed downregulation of core ubiquitin-coding genes. In contrast, deubiquitinating enzyme genes showed minimal or no significant change, indicating selective disruption of ubiquitin-associated regulatory pathways. (G) Expression analysis of ribosomal protein S3 (RPS3) in astronaut skin biopsies across epidermal, dermal, and vascular compartments from four astronaut subjects. RPS3 gene levels were reduced post-flight compared to pre-flight samples. (H–I) Expression of RPS3 in DA system. (H) DA organoids exposed to simulated microgravity (SMG) versus 1G controls with reduced RPS3 protein expression in SMG conditions. (I) Analysis of adhered differentiated SH-SY5Y DA neurons under SMG and 1G conditions confirmed downregulation of RPS3 protein expression under SMG. β-tubulin was used as a loading control. (N=3-5) (J) Global protein synthesis was assessed using the SUnSET assay, measuring puromycin incorporation. Results demonstrate a significant reduction in nascent protein synthesis in SMG-exposed cells compared to 1G controls, indicating impaired translational activity. All experimental data represent mean ± SEM followed by two-tailed unpaired Student’s t-test, *p < 0.05 considered statistically significant.

While protein degradation pathways, particularly the ubiquitin–proteasome system, have been extensively studied in PD, early steps of protein synthesis—especially ribosomal dysfunction— remain relatively underexplored, with only a few studies addressing this aspect. In contrast to the classical emphasis on protein aggregation and degradation failure, our data reveal a marked reduction in ribosomal pathway activity in the blood of PD patients.

We then analyzed transcriptomic data from peripheral blood mononuclear cells (PBMCs) of astronauts^32^, comparing post-flight samples to pre-flight baselines. We observed a robust and persistent downregulation of cytosolic ribosomal gene sets post-flight, which remained evident even up to 82 days after return (Fig 5E).

Next, we examined genes associated with the ubiquitin–proteasome system in astronaut blood. Interestingly, core ubiquitin-coding genes were downregulated, whereas deubiquitinating enzymes were not significantly affected (Fig 5F). Notably, certain ubiquitin genes such as RPS27A encode both ubiquitin and ribosomal proteins, suggesting a functional intersection between ribosomal and proteasomal pathways (ref). In addition, we identified alterations in spliceosomal pathways in PD blood datasets, indicating that RNA splicing may represent another key regulatory process disrupted both in spaceflight and in PD.

To identify a potential proxy for ribosomal protein, we focused on RPS3, a ribosomal protein for which reliable commercial antibodies are available. We assessed its expression in astronaut skin—an ectoderm-derived tissue similar in origin to the nervous system—and found that RPS3 protein levels were reduced post-flight (Fig 5G).

We next investigated whether microgravity directly affects protein translation in the DA system. Using DA organoids exposed to simulated microgravity, we observed a significant downregulation of RPS3 expression compared to 1G controls (Fig 5H). Similarly, Western blot analysis of differentiated SH-SY5Y DA neurons under simulated microgravity conditions revealed a marked decrease in RPS3 protein levels, consistent with a global downregulation of ribosomal proteins and pathways (Fig 5 I, J).

To determine whether these changes translate into functional deficits in protein synthesis, we performed a SUnSET assay to measure global translation. We observed a significant reduction in puromycin incorporation in the simulated microgravity group compared to 1G controls, indicating decreased nascent protein synthesis (Fig 5K).

Together, these findings demonstrate that spaceflight and microgravity not only impair mitochondrial function but also disrupt the global protein synthesis machinery. This effect is observed in blood as well as in DA systems, suggesting that microgravity exposure induces physiological changes that recapitulate key molecular features of PD.

### Post-SMG cell free DNA increase is specific to mitochondrial source and not nuclear across both adherent and organoid DA system

Elevated levels of cell free DNA (cfDNA) were detected in the culture supernatants of SMG exposed neurons compared to static 1G controls, indicating that simulated microgravity induces measurable cellular stress and may trigger damage associated molecular pattern (DAMP) signaling pathways (Fig 6). To determine the origin of this cfDNA, we quantified mitochondrial DNA (mtDNA; mtND5) and nuclear DNA (Hprt1) using multiplex digital PCR. SMG exposure resulted in a preferential increase in mtDNA derived cfDNA, with mtND5 levels significantly elevated relative to nuclear cfDNA. This selective enrichment of mitochondrial sequences suggests that mitochondria are the primary source of cfDNA release under SMG, rather than generalized nuclear breakdown.

**Figure 6.**
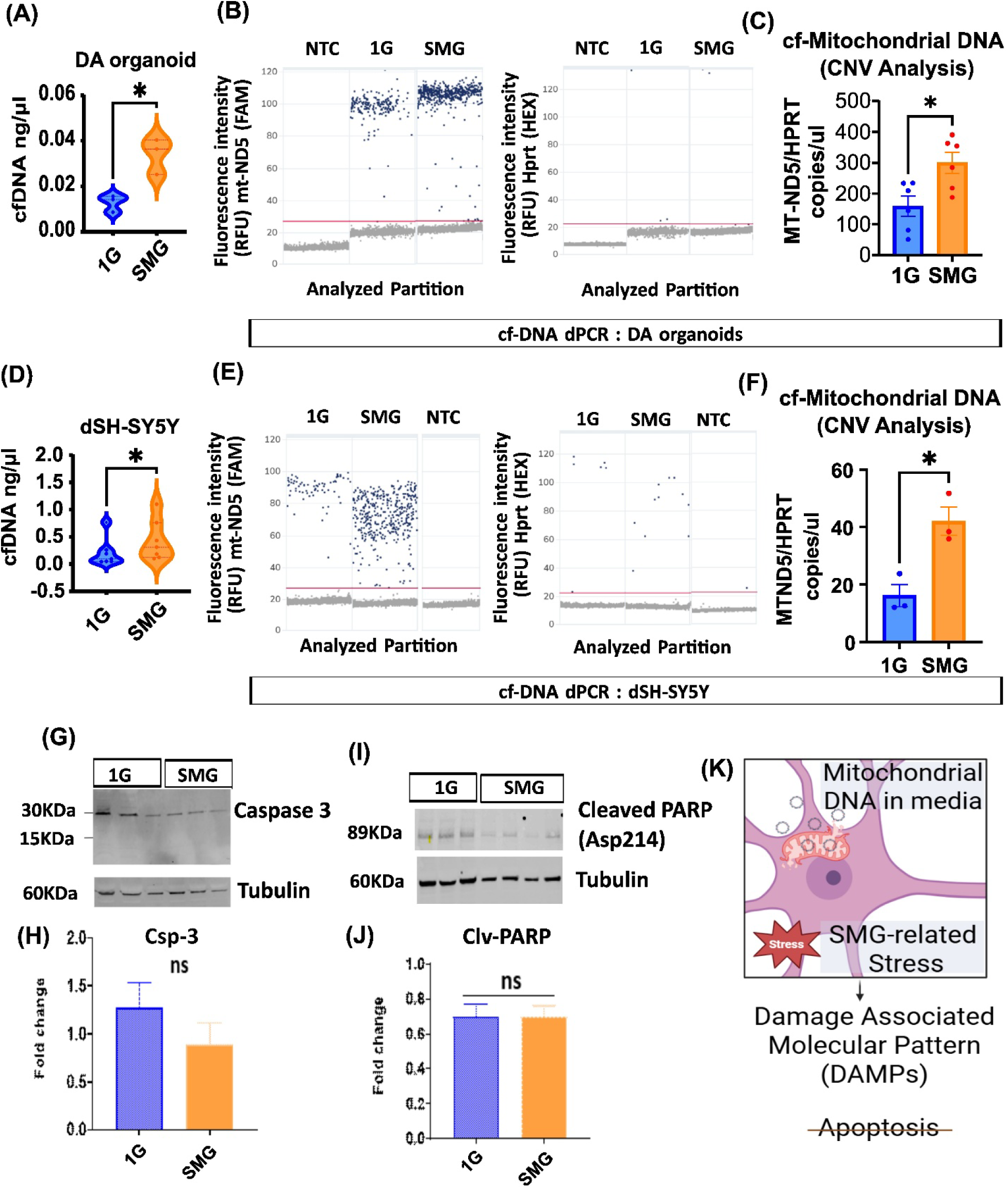
Simulated Microgravity (SMG) induces mitochondrial DNA release in organoids and dSH-SY5Y neurons without triggering apoptosis. (A) Representative analysis showing increased cell-free DNA (cfDNA) release in the culture media of DA organoids under simulated microgravity (SMG) conditions compared to 1G controls (n=3-5). (B) Multiplex digital PCR (dPCR) for the quantification of mitochondrial DNA (mtDNA) and nuclear DNA (nDNA). mtDNA is detected using mtND5 primers (FAM channel), and nDNA is detected using HPRT primers (HEX channel). (C) Copy Number Variation (CNV) analysis of released mtDNA in DA organoid media, demonstrating a significant increase in mtDNA release under SMG conditions. (n=6) (D) Analysis of cfDNA release in differentiated SH-SY5Y neuronal cultures under SMG vs. 1G conditions. (n=6) (E) Multiplex dPCR analysis of SH-SY5Y media confirming increased levels of ND5 (FAM) relative to HPRT (HEX) in SMG samples. (F) CNV analysis of cf-mtDNA in SH-SY5Y media, showing elevated copies per microliter under SMG (n=3). (G) Western blot analysis of SH-SY5Y cell lysates evaluating apoptosis markers Caspase-3 and Tubulin (loading control). No significant difference in the expression of cleaved caspase-3 (15 kDa) versus full-length caspase-3 (30 kDa) was observed between 1G and SMG groups. (n=3-6). (H) Densitometry quantification of caspase-3 expression, showing no significant apoptotic induction. (I) Western blot analysis of cleaved PARP (Asp214) expression normalized to tubulin (n=3-6) (J) Densitometry quantification of cleaved PARP, confirming no significant increase in apoptotic signaling. (K) Schematic representation illustrating that SMG-induced stress triggers the release of mtDNA, which may function as a Damage-Associated Molecular Pattern (DAMP), occurring independently of apoptotic cell death pathways. All experimental data represent mean ± SEM followed by two-tailed unpaired Student’s t-test, *p < 0.05 considered statistically significant.

Importantly, classical apoptotic markers—including cleaved PARP1 and cleaved caspase 3— remained unchanged (Fig. 6 G–J). The absence of apoptotic activation indicates that cfDNA release is not a consequence of cell death, but instead reflects sublethal mitochondrial stress, impaired mitochondrial quality control, or early senescence like phenotypes. Extracellular mitochondrial DNA has been reported to function as a damage-associated molecular pattern (DAMP) capable of activating innate immune pathways, including TLR9, cGAS–STING, and the NLRP3 inflammasome, in appropriate biological contexts. Thus, the selective increase in extracellular mtDNA under SMG conditions indicates mitochondrial stress in the absence of overt apoptotic cell death. These findings are consistent with mitochondrial stress-associated mtDNA release and raise the possibility that extracellular mtDNA could participate in downstream innate immune signaling, although the underlying release mechanism and functional consequences were not investigated in the present study.

### Proteomics data suggests SMG suppresses immune surveillance, splicing, proteostasis, mitochondrial, and DA signaling pathways

Proteomic profiling of differentiated SH-SY5Y cells exposed to simulated microgravity (SMG) revealed extensive remodeling of the cellular proteome compared with 1G controls. Volcano plot analysis identified a broad set of significantly upregulated and downregulated proteins, indicating a strong global response to altered gravitational conditions (Figure 7A). Network-based clustering of downregulated proteins identified two principal functional modules (Figure 7B). The dominant downregulated cluster comprised RNA processing and transcriptional regulatory proteins, including THRAP3, BCLAF1, and RSRC2, suggesting suppression of key components of mRNA splicing and transcriptional regulation under SMG (Figure 7C). In contrast, upregulated proteins formed multiple distinct functional clusters (Figure 8D). The largest cluster was enriched for apolipoproteins, lipid transporters, and secreted inflammatory or extracellular proteins, including APOE, APOA4, APOC3, APOM, APOO, PON1, PPARD, SAA4, SCN2A, JCHAIN, GPX3, SERPINF1/2, THBS1, AHSG, ITIH2, ITIH3, and AFP, indicating a strong shift toward lipid metabolism, extracellular signaling, and secretory pathway activation under SMG conditions. Additional clusters included ERVMER34-1, MRS2, RBM5 (Cluster 2), spliceosome-associated proteins CWC15 and CWC27 (Cluster 3), extracellular matrix proteins LUM, FMOD, and PRG4 (Cluster 4), and mitochondrial/oxidative phosphorylation-related proteins CLCC1, MTCO3, MT-ATP, and SUPV3L1 (Cluster 5). Several functionally relevant proteins, including IKBIP, BAD, AOC3, DHX29, SLC25A6, and THRB, remained unclustered but were significantly regulated (Figure 7E). Functional enrichment analysis further supported these observations. Upregulated proteins were significantly enriched in pathways related to amyloid-associated processes, HDL and lipoprotein particle remodeling, lipid metabolic disorders, and PPAR signaling (Figure 7F). In contrast, downregulated proteins were strongly associated with RNA metabolic processes, including mRNA processing, spliceosome organization, nuclear speck function, and RNA splicing pathways (Figure 7G). Together, these findings demonstrate that simulated microgravity induces a coordinated proteomic response in SH-SY5Y cells characterized by suppression of RNA processing machinery and activation of lipid metabolism, extracellular secretion, and stress-associated signaling pathways.

**Figure 7.**
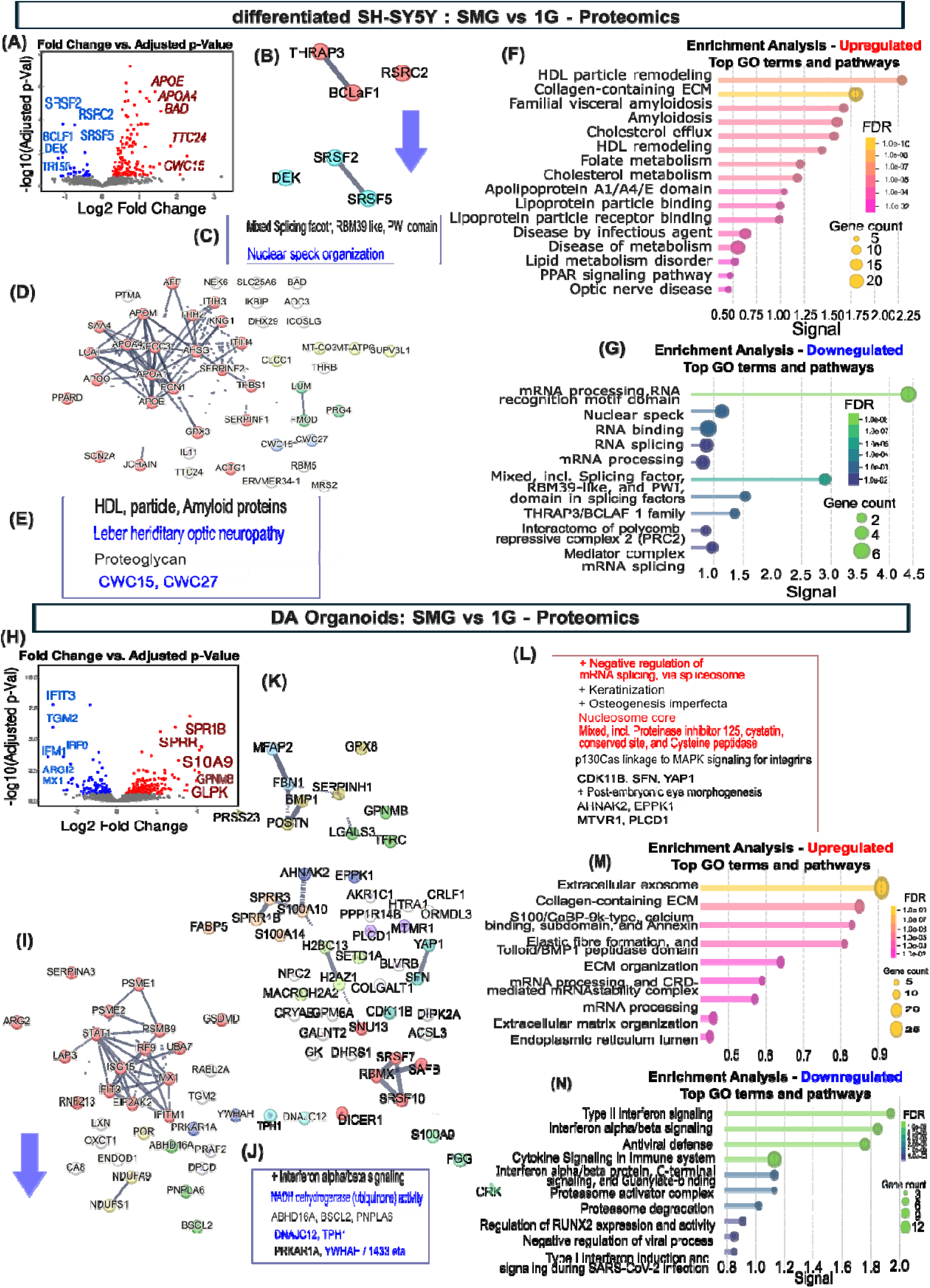
Protein-protein interaction and cluster analysis of Proteomics data from differentiated SH-SY5Y neurons and DA organoids exposed to SMG versus 1G conditions. (A) Volcano plot showing differentially expressed proteins in differentiated SH-SY5Y cells under SMG compared with 1G controls. Significantly upregulated and downregulated proteins are highlighted based on fold-change and adjusted p-value thresholds. (B) STRING-based network clustering of downregulated proteins identifying two primary functional modules. (C) Functional characterization of downregulated clusters. Cluster 1 consists of RNA processing and transcription-associated factors including THRAP3, BCLAF1, and RSRC2, indicating suppression of RNA splicing and transcriptional regulatory machinery under SMG. (D) STRING-based network clustering of upregulated proteins revealing multiple distinct functional modules. (E) Functional annotation of upregulated protein clusters. Cluster 1 is enriched for lipid transport, apolipoproteins, and inflammatory/secreted proteins including APOM, SAA4, APOO, APOE, PON1, PPARD, SCN2A, JCHAIN, GPX3, SERPINF1/2, THBS1, AHSG, ITIH2, ITIH3, AFP, and APOA4. Additional clusters include: Cluster 2 (ERVMER34-1, MRS2, RBM5), Cluster 3 (CWC15, CWC27), Cluster 4 (LUM, FMOD, PRG4), and Cluster 5 (CLCC1, MTCO3, MT-ATP, SUPV3L1). Unclustered upregulated proteins include IKBIP, BAD, AOC3, DHX29, SLC25A6, and THRB. (F) Gene Ontology (GO) enrichment analysis of upregulated proteins showing significant enrichment of pathways related to amyloid-associated processes, HDL/lipoprotein particle remodeling, lipid metabolism disorders, and PPAR signaling. Top enriched biological processes and pathways are shown together with false discovery rate (FDR) values and gene counts. (G) GO enrichment analysis of downregulated proteins highlighting significant suppression of pathways involved in mRNA processing, spliceosome function, nuclear speck organization, and RNA splicing. (H) Volcano plot of differentially expressed proteins in DA organoids cultured under SMG versus 1G conditions, showing fold change versus adjusted P value and highlighting significantly altered proteins. (I) STRING network analysis of downregulated proteins in DA organoids exposed to SMG. (J) Functional annotation of downregulated clusters in DA organoids, including interferon-α/β signaling, mitochondrial NADH dehydrogenase-associated proteins, DA signaling-associated proteins DNAJC12, TPH1, and a 14-3-3 zeta (YWHAZ)-associated cluster. (K) STRING network analysis of proteins upregulated in DA organoids following SMG exposure. (L) Functional characterization of upregulated clusters, including proteins involved in negative regulation of mRNA splicing, spliceosome-associated factors, nucleosome core components, protease inhibitors, cysteine peptidase regulators, and clusters containing YAP1-associated proteins. (M) GO and pathway enrichment analysis of proteins upregulated in DA organoids under SMG. Extracellular exosome-related pathways, collagen and extracellular matrix (ECM) secretion, ECM organization, and mRNA processing pathways are significantly enriched. (N) GO and pathway enrichment analysis of downregulated proteins in DA organoids under SMG. Significantly suppressed pathways include interferon signaling, cytokine signaling, antiviral defense responses, proteasome-mediated degradation, and negative regulation of viral processes.

**Figure 8.**
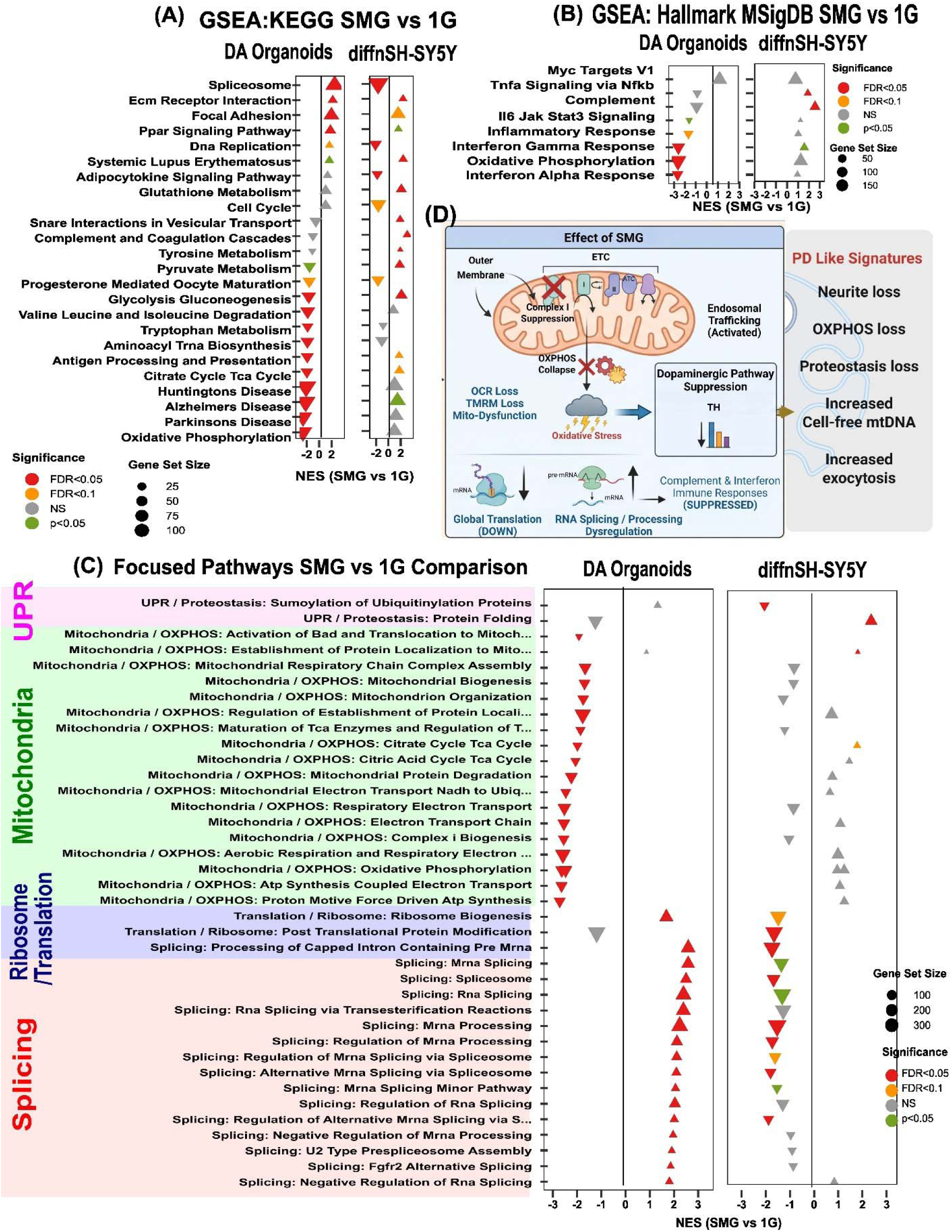
Gene Set Enrichment Analysis (GSEA) of Proteomics Data from Differentiated SH-SY5Y Neurons and DA Organoids Exposed to SMG versus 1G Conditions. (A–B) Unbiased GSEA of simulated microgravity (SMG) vs. 1G controls in both DA organoids and differentiated (diff.) SH-SY5Y neurons, displaying enriched pathways on the y-axis and normalized enrichment scores (NES) on the x-axis for (A) KEGG Legacy Pathways and (B) Hallmark Molecular Signatures Database (MSigDB) pathways. (C) Detailed, focused analysis of key enriched pathways—including the Unfolded Protein Response (UPR), mitochondrial pathways, ribosomal/translation pathways, and splicing pathways—demonstrating divergent directionality between the 3D organoid system and 2D adherent neurons. For all plots (A–C), triangle-lollipop size corresponds to the gene set size. Color denotes statistical significance: red (FDR < 0.05), orange (FDR < 0.1), green (p-value < 0.05), and gray (not significant). Downward and upward arrows indicate negatively and positively enriched pathways, respectively. (D) Schematic representation illustrating the key impacts of SMG on the DA system, highlighting mitochondrial dysfunction, oxidative stress, altered translation and splicing pathways, and increased endosomal trafficking and exocytosis. These cellular disruptions collectively culminate in early PD-like signatures, including neurite loss, neuronal stress, and the downregulation of DA markers.

Having established the baseline stress response in 2D cultures, we next profiled DA organoids to determine how a 3D tissue architecture influences this adaptation. Proteomic profiling of these organoids cultured under SMG revealed extensive remodeling compared with 1G controls (Fig. 7H), exposing a distinct, highly coordinated response. STRING network analysis identified several functionally distinct clusters of downregulated proteins (Fig 7 I, J). The largest downregulated cluster consisted predominantly of proteins involved in Ubiquitin Proteasome pathways PSME1, PSME2, UBA7; Global translation regulator EIF2AK2, interferon signaling, antiviral defense, and innate immune response proteins STAT1, IFITM1, MX1, GSDMD, IRF9, RNF213, ARG2, and SERPINA3, indicating a marked suppression of immune and antiviral signaling pathways under SMG conditions. A second cluster contained mitochondrial respiratory chain components, including NDUFS1, NDUFA9, and POR, suggesting impaired mitochondrial electron transport and oxidative metabolism. Additional downregulated modules included proteins involved in lipid metabolism and membrane homeostasis (BSCL2, PNPLA6, ABHD16A), neuronal signaling proteins (PRKAR1A, YWHAH), and DA neuron-associated proteins (TPH1 and DNAJC12) (Fig 7 I, J). Consistent with these network findings, pathway enrichment analysis demonstrated significant downregulation of immune-related pathways, including interferon signaling, cytokine signaling, antiviral defense mechanisms, proteasome-mediated degradation, and negative regulation of viral processes (Fig 7). Collectively, these results indicate that SMG suppresses innate immune signaling, mitochondrial function, lipid homeostasis, and key neuronal pathways in DA organoids.

STRING network analysis of proteins upregulated in DA organoids under SMG revealed multiple interconnected functional modules (Figure 7K–L). The largest cluster was enriched for RNA-processing and splicing factors, including SNU13, SRSF7, SAFB, SRSF10, and DICER1, indicating activation of post-transcriptional regulatory mechanisms. A second cluster containing S100A9, FGG, and CRK suggested enhanced inflammatory and cell-signaling processes. Additional upregulated clusters included proteins involved in cell-cycle regulation and Hippo signaling (CDK11B, SFN, YAP1), chromatin organization and epigenetic regulation (H2AZ1, MACROH2A2, H2BC13, SETD1A), and epithelial differentiation-associated proteins (FABP5, SPRR1B, SPRR3, S100A10). Notably, several clusters were strongly associated with extracellular matrix (ECM) organization and remodeling. These included structural ECM proteins (MFAP2, FBN1) as well as matrix-remodeling and secretory proteins (PRSS23, POSTN, BMP1, SERPINH1, GPX8). An additional signaling-associated cluster containing PLCD1 and MTMR1 was also significantly upregulated. Consistent with these network observations, functional enrichment analysis demonstrated significant upregulation of extracellular exosome pathways, extracellular matrix organization, collagen-related pathways, ECM secretion, and mRNA-processing functions (Figure 7M). Together, these findings suggest that SMG promotes activation of RNA-processing programs, chromatin remodeling, extracellular matrix deposition and remodeling, and YAP1-associated mechano-transduction pathways in DA organoids. One particularly interesting biological interpretation is that the simultaneous upregulation of YAP1, POSTN, FBN1, MFAP2, BMP1, and collagen/ECM pathways points toward a coordinated mechano-adaptive extracellular matrix remodeling response to simulated microgravity, while the downregulation of STAT1/IRF9/MX1/IFITM1 indicates suppression of interferon-driven immune signaling.

### Global pathway analysis reveals fundamentally distinct simulated microgravity responses in 3D DA organoids and SH-SY5Y neurons

We performed preranked Gene Set Enrichment Analysis (GSEA) using t-statistic-ranked proteins across the Hallmark, KEGG, Reactome, and Gene Ontology Biological Process collections (MSigDB). Following gene symbol mapping, 4,602 proteins from DA organoids and 4,412 proteins from SH-SY5Y cells were analyzed, evaluating 3,698 and 3,606 gene sets, respectively, using 10,000 permutations (minimum gene set size = 15, maximum = 500; Benjamini-Hochberg FDR correction). Consistent with the protein-protein interaction networks identified by STRING, pathway-level analysis demonstrated that simulated microgravity elicited markedly different biological responses in the two experimental systems (Fig. 8). Across all MSigDB collections, 294 significantly enriched pathways exhibited opposite regulation between organoids and SH-SY5Y cells, whereas only 21 pathways were concordantly regulated (FDR < 0.05), indicating that cellular architecture profoundly influences the molecular response to altered gravitational loading.

Hallmark pathway analysis revealed a pronounced suppression of oxidative phosphorylation, interferon-α and interferon-γ responses, and inflammatory signaling in DA organoids following simulated microgravity exposure (Fig. 8A). In contrast, differentiated SH-SY5Y cells preferentially activated complement signaling, TNFα/NFκB-mediated inflammation, and additional stress-response pathways, consistent with a generalized inflammatory adaptation rather than a neuron-specific degenerative program.

KEGG pathway analysis further highlighted the divergence between the two models (Fig. 8B). DA organoids exhibited coordinated suppression of oxidative phosphorylation together with multiple neurodegeneration-associated pathways, including Parkinson’s disease, Alzheimer’s disease, and Huntington’s disease, accompanied by significant activation of spliceosomal pathways. Conversely, SH-SY5Y cells preferentially enriched complement and coagulation cascades, extracellular matrix remodeling, and related signaling programs while suppressing DNA replication. Notably, glycolytic metabolism and spliceosomal pathways displayed opposite enrichment patterns between the two models, underscoring fundamentally different adaptive responses to simulated microgravity.

Focused analysis of pathways associated with neurodegeneration further emphasized the distinct biological programs engaged in each model (Fig. 8C). DA organoids demonstrated coordinated activation of RNA processing and spliceosomal pathways, with all 16 significantly enriched splicing-related pathways increased following simulated microgravity exposure. This was accompanied by widespread suppression of mitochondrial respiration and oxidative phosphorylation, with 19 mitochondrial pathways significantly downregulated, together with reduced ribosome biogenesis and translational pathways. In contrast, these mitochondrial programs were largely absent in SH-SY5Y cells, which instead preferentially activated unfolded protein response and protein-folding pathways, indicative of a canonical cellular stress response. Despite these extensive differences, a limited set of biological processes was consistently regulated in both systems. The 21 concordantly enriched pathways were predominantly associated with extracellular matrix organization, cell adhesion, tissue remodeling, and wound-healing processes (Fig. 8D), suggesting that mechanotransduction-dependent remodeling represents a conserved response to simulated microgravity independent of neuronal maturity or three-dimensional tissue organization^30^.

Collectively, these findings demonstrate that while both experimental systems respond to altered gravitational loading, human DA organoids more faithfully recapitulate the coordinated mitochondrial suppression, translational dysfunction, and RNA-processing alterations characteristic of PD. In contrast, SH-SY5Y cells also exhibit overlapping SMG-associated molecular features, including dysregulation of proteostasis and immune-related pathways, increased exocytosis and secretory signaling programs, and loss of spliceosome-associated regulation, consistent with a partial convergence toward PD-like stress signatures. However, these responses are more fragmented and lack the integrated mitochondrial–translational axis observed in DA organoids, underscoring the superior fidelity of three-dimensional human DA models in capturing disease-relevant mechanisms induced by simulated microgravity.

### Bulk RNA-seq Analysis of the PPMI FOUNDIN-PD Cohort Reveals Early Translational and Mitochondrial Dysfunction in iPSC-Derived DA Neurons from PD patients

To determine whether pathways identified in our simulated microgravity (SMG) model are also altered in human PD, we analyzed bulk RNA-sequencing datasets from the Michael J. Fox Foundation (MJFF) Parkinson’s Progression Markers Initiative (PPMI) FOUNDIN PD dataset.

The dataset comprised iPSC-derived DA neurons generated from healthy controls and individuals with PD representing diverse genetic backgrounds, including idiopathic PD, GBA-positive PD, LRRK2-positive PD, GBA/LRRK2 comorbid cases, prodromal subjects, genetically unaffected carriers, and other PD-related genetic subgroups, however for our study we narrowed the analysis to idiopathic PD vs healthy control. Neuronal at different differentiation stages were assessed at day 0, day 25, and day 65, corresponding to neural induction, immature DA neurons, and mature DA neurons, respectively.

Gene Set Variation Analysis (GSVA) was performed across all patient samples using Hallmark Msigdb pathways followed by custom-curated gene sets showing highly significant alteration. Individual columns represent individual patient-derived neuronal cultures, while rows represent pathway enrichment scores (Fig 9 A). Multiple pathways previously implicated in PD and identified in our SMG studies exhibited coordinated alterations across disease groups. Prominent changes were observed in pathways associated with protein synthesis, mitochondrial bioenergetics, RNA processing and proteostasis.

**Figure 9.**
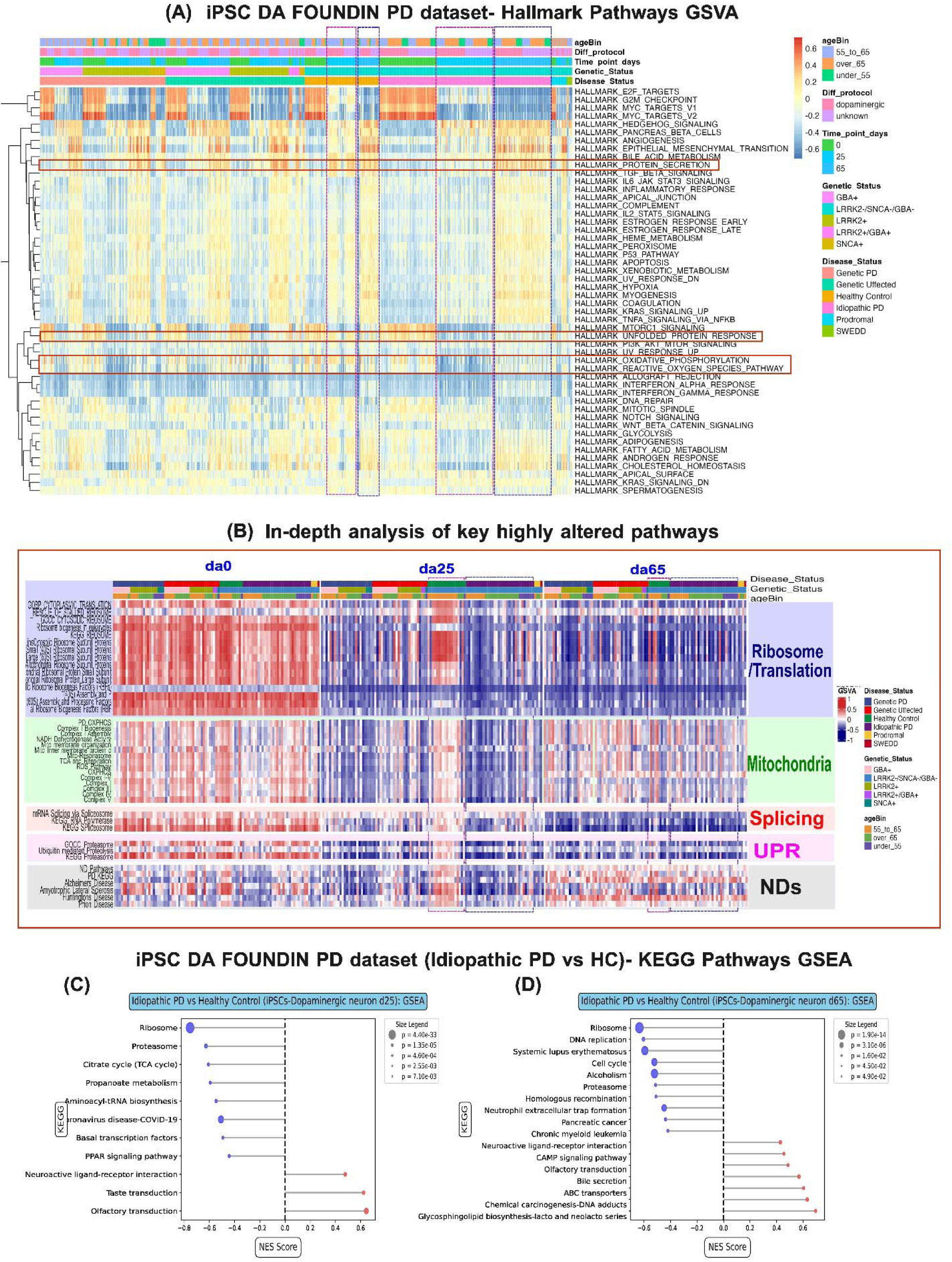
Transcriptomic analysis of PPMI-derived iPSC DA neurons identify ribosomal and mitochondrial dysfunction as a hallmark of Parkinson’s disease. (A) Gene Set Variation Analysis (GSVA) of bulk RNA-seq data from iPSC-derived DA neurons generated from the Parkinson’s Progression Markers Initiative (PPMI) cohort. Samples include healthy controls and PD subjects representing multiple genetic backgrounds, including idiopathic PD, GBA-associated PD, LRRK2-associated PD, GBA/LRRK2 comorbid cases, prodromal individuals, genetically unaffected carriers, and additional PD-related subgroups. Neurons were analyzed at day 0, day 25, and day 65 of differentiation. Each column represents an individual patient-derived neuronal sample, and each row represents a Hallmark pathway gene set. Red boxed regions highlight pathways significantly altered in PD and relevant to mechanisms identified in simulated microgravity studies. (B) Expanded GSVA using custom-curated pathway collections focused on Parkinson’s disease-relevant biological processes. Translation-associated pathways included KEGG ribosome, ribosome biogenesis, ribosomal protein components, and ribosome stalling gene sets. Mitochondrial pathways included curated electron transport chain (ETC) complexes I–V and oxidative phosphorylation signatures. Additional pathways analyzed included mRNA splicing and spliceosome function, unfolded protein response (UPR), ubiquitin-proteasome system activity, and neurodegenerative disease pathways derived from Parkinson’s disease, Alzheimer’s disease, Huntington’s disease, and prion disease gene sets. Pathway alterations were most pronounced at day 25 of differentiation. (C) Gene Set Enrichment Analysis (GSEA) comparing idiopathic PD and healthy control DA neurons at day 25 of differentiation. Enrichment results demonstrate significant dysregulation of ribosomal and translational pathways, accompanied by alterations in inflammatory signaling, proteostasis, and mitochondrial metabolism. (D) GSEA comparing idiopathic PD and healthy control DA neurons at day 65 of differentiation. Persistent suppression of ribosome-associated and protein synthesis pathways is observed in mature PD neurons, together with alterations in proteasome function, oxidative phosphorylation, and tricarboxylic acid (TCA) cycle pathways. These findings identify translational dysfunction as a conserved molecular signature of Parkinson’s disease DA neurons.

To further investigate these observations, expanded GSVA was performed using custom-curated pathway collections (Fig 9 B). Translation-related signatures included KEGG ribosome pathways, ribosomal protein genes, ribosome biogenesis pathways, and ribosome stalling/translation regulation gene sets. Mitochondrial pathways included curated electron transport chain (ETC) complexes I–V and oxidative phosphorylation genes derived from PD-focused datasets. Additional analyses examined mRNA splicing and spliceosome-associated pathways, unfolded protein response (UPR), ubiquitin-proteasome system components, and neurodegenerative disease pathways encompassin, Alzheimer’s disease, Huntington’s disease, and prion disease signatures.

Interestingly, pathway alterations were most pronounced at day 25 of DA differentiation, suggesting that early neuronal maturation stages may represent a critical window during which disease-associated molecular defects emerge. In contrast, day 0 cells displayed relatively limited differences, while many alterations persisted into day 65 mature neurons. Given the focus of the current study on disease-related cellular mechanisms relevant to simulated microgravity, subsequent analyses concentrated primarily on comparisons between healthy control and idiopathic PD neurons.

To validate the GSVA findings, Gene Set Enrichment Analysis (GSEA) for KEGG pathways was performed comparing idiopathic PD and healthy control neurons at day 25 and day 65 (Fig 9 C, D). Consistent across both developmental stages, translational and ribosomal pathways emerged among the most significantly downregulated biological processes in PD neurons. Additional dysregulated pathways included inflammatory signaling, proteasome function, mitochondrial metabolism, oxidative phosphorylation, and tricarboxylic acid (TCA) cycle pathways.

Collectively, these analyses identify impaired protein synthesis, mitochondrial dysfunction, and disruption of proteostasis as prominent and reproducible molecular signatures of PD DA neurons. The remarkable overlap of these pathways with those altered under simulated microgravity supports the hypothesis that SMG recapitulates core molecular mechanisms associated with early PD pathogenesis and provides a tractable experimental platform for studying disease initiation and progression.

## Discussion

To our knowledge, this is the first study to interrogate human iPSC-derived DA midbrain organoids and 2D differentiated SH-SY5Y DA neurons under simulated microgravity (SMG) and to define the cellular and molecular response. Using a 2D clinostat calibrated to deliver acceleration vectors approaching deep-space transit conditions (∼0.048 G at 15 rpm), we show that 72 h of SMG is sufficient to elicit a coordinated, sublethal stress state across structural, functional and molecular axes that converges on hallmarks of early PD: early neurite retraction, DA identity loss, engagement of familial-PD mitochondrial quality-control kinases, mitochondrial bioenergetic failure, translational repression, and selective extraceullar release of mitochondrial DNA in the absence of apoptotic activation. Unlike toxin-based or monogenic PD paradigms, which precipitate neuronal death within hours to days, SMG produces a multi-system perturbation that captures an early, potentially reversible window of DA vulnerability.

The earliest morphological readout of SMG was reduced neurite length in 2D DA neruons and reduced organoid size and sprouting complexity in 3D midbrain organoids, with neurite width preserved. This indicates selective impairment of process extension rather than gross cytoskeletal collapse. Cytotoxicity by LDH release was modest (∼10%), arguing against widespread cell death as the driver. Crucially, gene set enrichment analysis of FOUNDIN-PD iPSC-derived DA neuron datasets^33^ revealed downregulation of cytoskeletal regulation, cell-membrane projection, neurite, and tunneling-nanotube pathways in both prodromal and idiopathic PD relative to controls, while axonogenesis pathways were paradoxically upregulated, which is consistent with compensatory neurite-growth signaling against a backdrop of failing structural maintenance. The convergence between SMG-exposed organoids and prodromal PD transcriptomes positions early process retraction as a shared, possibly upstream feature of DA vulnerability, predating frank neuronal loss.

At the molecular level, SMG produced a transcriptional signature reminiscent of familial PD: downregulation of *TH* and *SNCA* alongside upregulation of *PINK1*, *PRKN*, and *LRRK2*. Tyrosine hydroxylase is the rate-limiting enzyme of dopamine biosynthesis and an early casualty of DA stress in human PD substantia nigra and in iPSC-based models^34^. Consistent with this loss of DA identity, organoid proteomic profiling revealed downregulation of DNAJC12, a co-chaperone required for stability of the catecholaminergic biosynthetic machinery, further supporting disruption of monoaminergic enzymatic programs and suggesting that impaired proteostasis may precede or accompany TH depletion. Reduction at both transcript and protein levels indicates erosion of DA identity rather than transient regulatory shift. The coordinated upregulation of the canonical mitophagy axis, *PINK1* and *PRKN*^34^, together with *LRRK2*, signals engagement of the mitochondrial quality-control machinery whose dysfunction underlies a substantial fraction of monogenic PD; pathogenic *LRRK2* in particular interferes with PINK1/Parkin-dependent mitophagy and promotes mitochondrial aggregation^35,36^. Notably, extracellular dopamine concentrations remained unchanged despite this transcriptional, translational and electrophysiological compromise, suggesting transient compensatory preservation of neurotransmitter output (possibly by increased vesicular release efficiency, mobilization of reserve pools, or reduced reuptake) that masks the underlying degenerative trajectory. This dissociation mirrors the prodromal phase of human PD, in which substantial nigrostriatal injury accumulates before motor symptoms emerge^37^.

Mitochondrial dysfunction emerged as the most pervasive and quantitatively robust feature of the SMG response, spanning bioenergetics, complex composition, organelle distribution, and DNA integrity. TMRM imaging revealed loss of mitochondrial membrane potential in both somatic and neuritic compartments of 2D DA neurons and in DA organoids, a readout widely associated with impaired electron transport chain function and increased susceptibility to cell stress and degeneration^38,39^. Axonal segments showed particularly pronounced depletion of TMRM-positive mitochondria, consistent with disrupted mitochondrial trafficking — a process essential for axonal maintenance and synaptic function and a known vulnerability point in DA neurons whose long, highly branched axonal arbors and elevated bioenergetic demand render them uniquely susceptible to mitochondrial perturbation^40–43^. Real-time oxygen consumption measurements showed that, while baseline respiration was comparable between conditions, SMG-exposed organoids failed to recover metabolic activity following replating, plateauing at approximately 8% of 1G respiratory output by 168 h. Immunoblotting revealed selective loss of OXPHOS Complexes I, II and IV with preservation of Complex III, mirroring the electron-transport chain deficits reported in PD substantia nigra^44,45^, and compensatory upregulation of Complex V consistent with attempted preservation of ATP synthesis under upstream chain failure^46^. Together, these data position mitochondrial impairment not as an accompaniment of SMG-induced DA stress but as its central upstream driver, integrating with the elevated *PINK1*/*PRKN* signal as evidence of an active but ultimately insufficient quality-control response.

Mitochondrial dysfunction was accompanied by a profound suppression of protein synthesis, revealing a second conserved axis of the SMG response that closely parallels both human PD and spaceflight biology. Re-analysis of the FOUNDIN-PD iPSC-derived DA neuron dataset demonstrated that ribosomal pathways, ribosome biogenesis, oxidative phosphorylation, mitochondrial metabolism, and RNA-processing, spliceosome pathways are among the earliest molecular alterations at early DA neurons. Similar translational signatures were observed in independent PDBP peripheral blood datasets, indicating that these abnormalities extend beyond the central nervous system. While PD research traditionally focuses on downstream protein aggregation and the unfolded protein response (UPR), upstream disruptions in RNA processing, splicing and protein synthesis have emerged as critical pathogenic drivers. Accumulating evidence highlights that spliceosome anomalies and subsequent RNA mis-splicing, generation of circular RNA disrupt cellular homeostasis^47–50^, while aberrant ribosomal translation and protein synthesis anomalies directly induce proteotoxic stress^51–54^, collectively accelerating DA neurodegeneration. Notably, astronaut peripheral blood mononuclear cell transcriptomes likewise exhibited persistent downregulation of cytosolic ribosomal pathways following long-duration spaceflight, suggesting that translational suppression represents a conserved response to gravitational stress^55,56^. Consistent with these transcriptomic observations, SUnSET assays demonstrated a functional reduction in global protein synthesis under SMG, while RPS3 was independently reduced in astronaut skin, SMG-exposed DA organoids, and differentiated SH-SY5Y neurons. Mitochondrial dysfunction and translational repression are mechanistically coupled (protein synthesis is among the most ATP-demanding cellular processes, and the integrated stress response coordinately suppresses translation under bioenergetic strain)^57–59^ suggesting a feed-forward loop in which mitochondrial failure constrains translation and translational failure further compromises mitochondrial protein renewal. This translationally repressed state may also explain why *SNCA* downregulation occurs without overt α-synuclein accumulation in this 72h window, capturing a pre-aggregation phase of DA stress not modelled by existing toxin or overexpression paradigms. Together, these findings define a tightly coupled mitochondrial–translational stress network that converges on progressive destabilization of DA identity, recapitulating early, pre-degenerative features of PD across independent human datasets and experimental platforms.

A defining feature of the SMG phenotype is its sublethal character. Despite extensive mitochondrial, transcriptional and translational perturbation, classical apoptotic markers (cleaved PARP1 and cleaved caspase-3) were unchanged, and cytotoxicity was modest. Nonetheless, cell-free DNA in conditioned media was significantly elevated, and digital PCR demonstrated that this increase was selectively driven by mitochondrial DNA (*MT-ND5*) rather than nuclear DNA (*HPRT1*). Mitochondrial DNA carries CpG-rich, hypomethylated, bacterial-derived motifs that engage TLR9^60^, cGAS–STING^61^, and NLRP3 to amplify sterile inflammation, and circulating mtDNA has been implicated in age-related and neurodegenerative inflammatory priming. Selective extracellular mtDNA release in the absence of apoptotic signaling therefore points to sublethal mitochondrial membrane permeabilization or active mtDNA extrusion (possibly via VDAC oligomers or mitochondrial-derived vesicles) rather than terminal cellular dissolution. This signature distinguishes the SMG model from acute death-driven systems and suggests that microgravity may prime DA neurons for chronic neuroinflammatory crosstalk, a mechanism increasingly recognized as a contributor to DA vulnerability in PD^62^. A recent report suggest selective mitoDNA and neurofilamnet release from PD mid brain organoids complementing our findings^63^.

Proteomic profiling provided mechanistic insight into the adaptive response induced by SMG and revealed a striking divergence between 2D neurons and 3D organoids. 3D DA organoids mount a coordinated mechano-adaptive response that is entirely absent in 2D SH-SY5Y cultures. In the 3D environment, gravitational unloading activates a structural remodeling program driven by the upregulation of the Hippo effector YAP1 and extracellular matrix proteins (POSTN, FBN1, MFAP2, BMP1). Crucially, this physical adaptation runs parallel to a profound cellular collapse that mirrors both spaceflight-induced immunosuppression and the pathological hallmarks of PD. This neurodegenerative signature is defined by the suppression of the STAT1/IRF9 innate immune axis, deactivated interferon-stimulated genes (MX1, IFITM1), and silenced pyroptotic machinery (GSDMD), occurring in tandem with deficits in the ubiquitin-proteasome system (PSME1/2, UBA7), mitochondrial complex I (NDUFS1, NDUFA9), and the EIF2AK2-mediated translation-brake pathways. Notably, this proteostatic decay coincides with the upregulation of human endogenous retrovirus ERVMER34-1, replicating the epigenetic derepression observed in advanced PD and astronauts. This vulnerable, stress-sensitized state is further underscored by the upregulation of the pro-apoptotic factor BAD without corresponding cleavage of executioner caspase-3 or PARP1 as observed in Western blot. This "primed but not executing" apoptotic profile indicates that while the organoids exhibit prolonged mitochondrial stress and progressive loss of DA identity markers (DNAJC12, TPH1), they remain in a pre-apoptotic window open to therapeutic rescue. Conversely, 2D monocultures bypass this structural and neuroimmune decay entirely, shifting their economy toward lipid transport (APOE, APOC3, APOM) and PPAR signaling. These findings underscore the necessity of 3D biomimetic tissue models to accurately capture how gravitational unloading acts as a multi-system environmental trigger converging on progressive neurodegenerative pathways.

One of the most important findings of this study is the remarkable convergence between human PD datasets and SMG-induced molecular changes. Independent transcriptomic analyses of FOUNDIN-PD ipsc derived DA neurons identified disruption of translation, mitochondrial metabolism, proteostasis, and RNA processing and splicing as early disease-associated signatures. These same biological processes emerged from unbiased proteomic GSEA analyses of SMG-exposed ipsc derived DA brain organoids despite differences in experimental platform and molecular readout. Rather than reproducing late-stage neurodegeneration characterized by overt neuronal loss and α-synuclein aggregation, SMG appears to model an earlier pathogenic state defined by bioenergetic stress, translational repression, mitochondrial quality-control activation, and progressive loss of neuronal identity. This convergence supports the utility of SMG as a complementary human model for investigating early mechanisms of PD pathogenesis.

Several limitations of this study merit consideration. Ground-based 2D clinostat rotation simulates the time-averaged cancellation of the gravity vector but does not fully recapitulate true microgravity, and our 72h exposure window captures early stress responses but cannot speak to the trajectory of long-term adaptation, recovery or progression. Pharmacological and mechanical rescue experiments, including mitochondrial-protective interventions and reload paradigms, will be essential to establish causality between mitochondrial reactive oxygen species, translational repression and the DA phenotype, and are the subject of ongoing work. The 2D SH-SY5Y system, while widely used, lacks the cellular diversity and three-dimensional architecture of the organoid model, and conversely the organoid system lacks vascularization, microglial input and the substantia nigra microenvironment of the adult human brain. Validation in spaceflown human DA cultures^14^ and complementary radiation-exposure paradigms will be needed to dissect the relative contribution of microgravity from other space stressors. Despite these caveats, the convergence we observe between SMG-exposed human DA neurons, PD transcriptomes, and astronaut data supports the biological coherence of the model.

Our findings provide the first direct experimental evidence in a human DA model supporting prior meta-analytic predictions that spaceflight-associated stressors converge on PD-relevant molecular pathways^7^. Rather than inducing acute neurotoxicity, simulated microgravity elicited a coordinated set of mitochondrial dysfunction, translational repression, and progressive loss of DA identity while preserving cell viability and extracellular dopamine output, consistent with an early, compensatory phase of neuronal dysfunction. These observations suggest that altered gravitational loading perturbs fundamental bioenergetic and mechanobiological processes that are sufficient to initiate PD-like molecular and functional phenotypes in the absence of PD neurotoxins or disease-causing genetic mutations.

Beyond their implications for human spaceflight, these findings establish simulated microgravity as a tractable, toxin-free, and non-genetic human model of early DA dysfunction. The convergence of mitochondrial impairment, suppression of ribosome-associated pathways, and proteostasis dysregulation across simulated microgravity, astronaut transcriptomic datasets, aging-associated molecular hallmarks, and independent PD datasets suggests that altered gravitational loading engages conserved stress responses that normally emerge during biological aging. These observations raise the possibility that simulated microgravity accelerates aspects of cellular aging in DA neurons, thereby exposing molecular vulnerabilities that precede overt neurodegeneration. Future studies should define the temporal limits of compensatory dopamine homeostasis, determine how altered mechanical loading is coupled to mitochondrial dysfunction, translational control, and α-synuclein proteostasis, and evaluate whether interventions that preserve mitochondrial function, restore mechanical loading, or support ribosomal activity can prevent progression toward irreversible neurodegeneration. Collectively, our data position gravitational biology as an emerging axis in neurodegeneration research and argue that the DA system may be uniquely sensitive to perturbations in mechanical and bioenergetic load^42^.

## Methods

### Cell culture and DA differentiation of SH-SY5Y neurons. –

The neuroblastoma cell line SH-SY5Y (ATCC, USA) was maintained in Dulbecco’s modified Eagle’s medium (DMEM/F12) supplemented with 15% fetal bovine serum (FBS; GIBCO, Invitrogen, USA, USA), 50 U/mL penicillin, 50 μg/mL streptomycin, and MEM-NEAA (Sigma-Aldrich, USA). Cells were maintained at 37 °C in a humidified incubator with 5% CO_2_. Differentiation into DA-like neurons was induced by treatment with 10 μM all-*trans* retinoic acid for 6 days.

### iPSC culture and DA organoid generation

Neural stem cells derived from the iPSC cell line AICS0013 (Coriell institute) were patterned via dual-SMAD inhibition and ventralized using sonic hedgehog (SHH), purmorphamine, and CHIR99021, followed by maturation in Neurobasal media supplemented with N2 and B27 and added BDNF, GDNF, ascorbic acid, db-cAMP, TGFβ3, and DAPT^3–6^. At day 25, midbrain progenitors were aggregated into three-dimensional DA organoids and cultured in low attachment U-bottom 96 well plate (Corning) for 10 days then transferred to maturation dishes or cryovials containing one organoid per cryovials and twice weekly media change.

### Simulated microgravity using a 2D clinostat

Simulated microgravity (SMG) was generated using a custom-built 2D clinostat device (SpinAxis BioSystem; SABS) designed to fit inside a standard cell culture incubator for prolonged culture. The instrument was calibrated to deliver a range of low-gravity environments simulating conditions from lunar to Martian gravity. Differentiated 2D SH-SY5Y neurons were plated into slide flasks and sealed with sterile rubber stoppers fitted with a needle to remove residual fluid and air bubbles, as previously described^17^. Sealed flasks were positioned at the center of the SABS clinostat and rotated at 15 RPM for 72 h (∼0.04 G) based on prior optimization with adherent cells^17^. Matched 1 G controls were prepared identically and maintained in the same incubator without rotation. DA organoids matured for one month were transferred into cryovials, sealed in the same manner, and either clino-rotated at 10 RPM or maintained at 1G for 72 h. Both 2D and 3D cultures were allowed to acclimatize for 24 h prior to harvesting and downstream analyses.

### Neurite outgrowth analysis

For 3D cultures, mature DA organoids (day 40 ± 5) were embedded in growth-factor–reduced Matrigel droplets or plated onto poly-D-lysine/laminin-coated glass-bottom dishes in defined differentiation medium to permit radial neurite extension, and were maintained for 48–96 h post-plating. For 2D differentiated SH-SY5Y cultures, phase-contrast images were calibrated to scale and converted to 8-bit format prior to analysis. Neurite length and branching were quantified in Fiji/ImageJ (v.1.54f) using the NeuronJ or Simple Neurite Tracer plugin. Measured parameters included total neurite length per organoid, average neurite length, and number of primary neurites. All analyses were performed in a blind manner from at least 12 fields per group 5 organoid per group and 7 independent fields per condition.

### Quantitative PCR

Total RNA was isolated using Trizol (Invitrogen, USA) method according to the manufacturer’s instructions, and concentration and purity of RNA were assessed by Qubit spectrophotometer (Invitrogen). Complementary DNA (cDNA) was synthesized from equal amounts of RNA using a reverse-transcription kit. Quantitative PCR was performed using SYBR Green chemistry on a LightCycler PCR machine (Roche). Gene expression was normalized to the housekeeping gene(s) [GAPDH], and relative expression was calculated using the 2^-ΔΔCt^ method.

### Digital PCR

Digital PCR (dPCR) was performed using the QIAcuity Digital PCR System (Qiagen) using custom primers and hydrolysis probes targeting mitochondrial NADH dehydrogenase subunit 5 (*MT-ND5*) and the nuclear reference gene hypoxanthine phosphoribosyltransferase 1 (*HPRT1*), designed via the GeneGlobe platform (Qiagen). The *MT-ND5* probe was labeled with FAM and the *HPRT1* probe with HEX to enable duplex detection. Reactions were prepared with QIAcuity dPCR Master Mix per the manufacturer’s instructions and loaded onto a QIAcuity Nanoplate 8.5k, partitioning each sample into approximately 8,500 nanowells for absolute quantification. Thermal cycling consisted of an initial denaturation at 95 °C for 2 min, followed by 40 cycles of 95 °C for 15 s and 60 °C for 30–60 s, with fluorescence acquisition in the FAM and HEX channels. Data were analyzed using QIAcuity Software (v. 3.1), with positive and negative partitions identified by fluorescence thresholds and target concentrations calculated using Poisson statistics. Results are reported as copies per microliter (copies/µL) of reaction.

### Dopamine quantification by LC-MS/MS

Dopamine levels in conditioned culture media were quantified by liquid chromatography-tandem mass spectrometry (MS). Media were mixed with trichloroacetic acid (TCA) at a 1:4 (v/v) ratio and incubated on ice for 10 min to precipitate proteins, then centrifuged at 14,000 × *g* for 15 min at 4 °C.The supernatant was collected for analysis, with identity confirmed by spiking with a DA standard, and quantification performed using the peak area under the curve. Analyses were carried out on an Agilent 1290 Infinity II LC system comprising a multisampler (G7167B), high-speed pump (G7120A), and multicolumn thermostat (G7116B), coupled to an Agilent 6470B triple quadrupole mass spectrometer. Chromatographic separation was performed on an Agilent InfinityLab Poroshell 120 EC-C18 analytical column (2.1 × 150 mm, 1.9 µm) preceded by a matching guard column (2.1 × 5 mm, 1.9 µm), with the column oven held at 40 ± 2 °C and the autosampler was held at 10 ± 2 °C. The injection volume was 2 µL and the total run time was 10 min. Mobile phases were 0.1% (v/v) formic acid in water (A) and 0.1% (v/v) formic acid in acetonitrile (B), Delivered at 0.25 mL min⁻¹ using the following gradient: 95% A/5% B at 0 min, 80% A/20% B at 2 min, 10% A/90% B at 2.1 min held until 10 min. Mass spectrometric detection used an Agilent Jet Stream electrospray ionization source in positive ion mode, with data acquired in multiple reaction monitoring (MRM) mode. Source parameters were: drying gas temperature, 250 °C; drying gas flow, 11 L min⁻¹; nebulizer pressure, 40 psi; sheath gas heater temperature, 400 °C; sheath gas flow, 12 L min⁻¹; capillary voltage, 4,000 V; and nozzle voltage, 0 V. Dopamine was monitored using the precursor-to-product ion transition *m/z* 154.1 to 137 with a 50 ms dwell time, fragmentor voltage of 131 V, collision energy to 8 V, and cell accelerator voltage of 5 V.

### Western Blotting

Cells were lysed in pre chilled RIPA buffer (Thermo Scientific) supplemented with phosphatase and protease inhibitor cocktail (Thermo Scientific) at a 100:1 (RIPA:PI) ratio. Lysates were incubated on ice for two 15 min intervals with intermittent vortexing and clarified by centrifugation at 13,000 × g for 15 min at 4 °C. Protein concentrations were determined using a BCA assay (Thermo Scientific). Equal amounts of protein were resolved by SDS–PAGE on 12% resolving / 4% stacking precast gels (Bio⍰Rad) at 35 mA per gel for 1 h and transferred to nitrocellulose membranes using semi⍰dry transfer. Membranes were blocked in Intercept (PBS) Blocking Buffer (LI⍰CORbio) for 1 h at room temperature, washed in PBS, and incubated overnight at 4 °C with primary antibodies. After three washes in TBS, membranes were incubated with the appropriate secondary antibodies (LI-CORbio IRDye 800CW, 700CW) for 1 h at room temperature, protected from light. Blots were washed in TBS and imaged on a LI COR Odyssey CLx Imaging System. LI⍰COR Chameleon Duo and Page Ruler Plus Stained Protein Ladder (Invitrogen, USA) were used as molecular weight markers.

### SUnSET puromycin incorporation assay

Global protein synthesis was assessed using the SUnSET assay. Cells were incubated with 10 µg/mL puromycin (Sigma Aldrich) for 15 min at 37 °C, washed with cold PBS, and lysed in RIPA buffer supplemented with protease inhibitors. Lysates were clarified by centrifugation at 14,000 × *g* for 15 min at 4 °C, and protein concentrations were determined by BCA assay. Equal amounts of protein were separated by SDS–PAGE and transferred onto nitrocellulose membranes. Membranes were blocked, incubated with anti-puromycin primary antibody (Sigma-Aldrich, #MABE343,), and probed with IRDye-conjugated secondary antibodies. Signals were detected and quantified on a LI-COR Odyssey imaging system.

### Whole-mount immunostaining, tissue clearing and confocal imaging of organoids

Organoids were fixed in 4% paraformaldehyde for 4 h at 4 °C and washed in PBS for 24 h. Samples were incubated with primary antibodies for 24 h at room temperature, washed five times for 10 min at room temperature, washed overnight at 4 °C. Secondary antibody incubation was performed for 24 h, followed by the same washing regimen. Tissue clearing was performed using RapiClear (Sunjin Lab), and organoids were mounted in RapiClear prior to imaging. Confocal images were acquired on a Leica STELLARIS confocal microscope.

### LDH cytotoxicity assay

Cytotoxicity was assessed using the CyQUANT™ LDH Cytotoxicity Assay kit (Invitrogen, USA) according to the manufacturer’s instructions. Cytotoxicity was measured using the CyQUANT™ LDH Cytotoxicity Assay (Invitrogen, USA) following the manufacturer’s protocol. Spontaneous LDH release, maximum LDH release, and experimental samples (50 µL media from cells cultured in 1G or SMG conditions) were collected, and maximum LDH was generated by a 45⍰min incubation at 37 °C with lysis buffer. Supernatants (50 µL) were transferred to a fresh 96⍰well plate, mixed with 50 µL Reaction Mixture, incubated for 10–30 min at room temperature protected from light, and stopped with 50 µL Stop Solution. Fluorescence (Ex 560 nm/Em 590 nm) was recorded, background⍰subtracted LDH activity was calculated, and percent cytotoxicity was determined as % cytotoxicity = [(experimental LDH − spontaneous LDH) / (maximum LDH − spontaneous LDH)] × 100

### Mitochondrial physiology and live imaging of membrane potential

Mitochondrial membrane potential was measured with the proton-gradient-sensitive dye TMRM (Invitrogen, USA, USA), following previously published protocol^7,8^. Cells were incubated with 20 nM TMRM for 30 min, at 37□C in 5% CO_2_ at 100% humidity. TMRM-stained cells were imaged in phenol red-free complete medium (GIBCO, USA) on a Leica STELLARIS 5 confocal microscope fitted with an Uno top-stage incubator maintaining 37□C and 5% CO_2_. Z-stack images were acquired at 40x magnification using a water-immersion objective, and maximum-intensity projections were generated in Fiji (v.1.54f).

### Cell-free DNA isolation from conditioned media

Cell-free DNA (cfDNA) was isolated from conditioned media using the DNeasy PowerViral Kit (Qiagen) according to the manufacturer’s instructions, with minor optimizations. Media collected from cultured cells or organoids were centrifuged sequentially at 300 × *g* for 5 min and 2,000 × *g* for 10 min at 4°C to remove cells and debris. Up to 500 µL of clarified supernatant was mixed with lysis buffer, incubated at room temperature for 10 min, and treated with inhibitor=removal reagents to eliminate PCR inhibitors. cfDNA concentration and fragment-size distribution were assessed using Qubit dsDNA High Sensitivity Assay (Invitrogen, USA).

### Real-time oxygen consumption rate measurements

Real-time oxygen consumption rate (OCR) was measured using the Resipher system (Lucid Scientific), a non-invasive, label-free metabolic monitoring platform that quantifies extracellular oxygen consumption via optical oxygen sensors positioned beneath each well of a standard multi-well plate, allowing continuous measurement of dissolved-oxygen depletion as a proxy for mitochondrial respiration dyes or media perturbation. Organoids were seeded into U-bottom (Corning) 96-well plates and monitored continuously over a 168-h experiment before and after either SMG or 1 G conditions. Following an initial on-plate growth phase (Pre SMG 0–45 h), organoids were removed and maintained off-plate under their respective gravity condition for 24 h, then re-seeded at ∼75 h for continued monitoring. Each condition included 12 organoid-containing wells and 4 cell-free control wells. OCR flux was recorded at 15-min intervals and expressed in fmol/mm²/s. Data were processed and analyzed using instrument software, with values averaged across wells and reported as Mean ± SEM. Periods corresponding to experimental interruptions (e.g., off-plate intervals or media changes) were excluded from direct measurement and are represented as interpolated regions in graphical outputs.

### Liquid Chromatography – Tandem Mass Spectrometry (LC-MS/MS) Based Proteomic Analysis

Twenty-five micrograms of total protein from each sample (n=3/group) were precipitated with cold acetone and redissolved in 8 M urea solution. After being reduced by dithiothreitol and alkylated by iodoacetamide, samples were diluted to 1M urea concentration and incubated with trypsin/LyC overnight at 30 °C. Digested samples were desalted and purified using a reverse-phase C18 spin column. Label-free quantitative proteomic analysis was performed on an Orbitrap Exploris 480 Mass Spectrometer (Thermo Fisher Scientific) coupled to a Vanquish Neo UHPLC system (Thermo Fisher Scientific) using a data independent acquisition (DIA) approach. Two micrograms of peptides were injected onto an EASY-Spray PepMap Neo reverse-phase C18 column (15cm x 75µm, 2 µm, 100 Å) and separated using a 65 min gradient with mobile phase A (0.1% formic acid in water) and B (80% acetonitrile, 0.1% formic acid in water) at a flow rate of 0.3 µl/min. Scan ranges for MS and MS/MS were set at 400 to 900 m/z with a resolution of 60,000 and 145 to 1,450 m/z with a resolution of 15,000, respectively. For DIA, 42 variable windows were used to cover m/z 400 to 900. Higher energy collisional dissociation (HCD) was set to 30% normalized collision energy (NCE) with default charge state set to +2. Acquired raw data was processed by Spectronaut (version 20.6, Biognosys, Newton, MA) using a library-free (directDIA) workflow with default settings and a protein sequence database for human downloaded from the UniProt.org on May 22, 2026. The FDR was set to 0.01 at the precursor and protein level.

### Bioinformatics Analysis

#### Gene Set Variation analysis (GSVA) FOUNDIN PD dataset analysis

Bulk RNA-sequencing data from iPSC-derived DA neurons from healthy controls and PD patients (n-98) were obtained from the FOUNDIN-PD initiative (PPMI, MJFF). Transcript count data derived from Salmon quantification were used to perform Gene Set Variance Analysis (GSVA) to calculate sample-wise pathway enrichment scores. A false discovery rate (FDR) threshold of 0.25 was used to identify significantly enriched pathways. GSVA package (v2.4.4) in R (v4.5.2) was used. Raw RNA-seq counts were assembled into a gene-by-sample count matrix using gene symbols as identifiers. Genes with a total count below 50 across all samples were removed, and counts were library-size normalized and variance-stabilized using the variance-stabilizing transformation (VST) in DESeq2 (v1.50.2) with an intercept-only design. Where multiple rows mapped to the same gene symbol, the entry with the highest mean expression across samples was retained. An overall GSVA using Hallmark MSigDB was used followed by in depth analysis of key pathways using custom curated Mitochondrial Electron transport chain and OXPHOS^7^, Proteostasis pathways including ribosome, translation, Proteasome, Ubiquitin mediated Proteolysis, Spliceosome, and Neurodegenerative diseases from KEGG. All pathways except custom curated ones are obtained from MSigDB. The resulting enrichment scores were visualized as a heatmap, where individual columns represent unique patient samples. GSVA was run on the variance-stabilized expression matrix using a Gaussian kernel for the cumulative density function estimation, with gene sets restricted to those containing between 15 and 500 detected genes and the maximum-deviation statistic used to define per-sample enrichment scores. This yielded a continuous sample-by-pathway score matrix reflecting the coordinated up- or down-regulation of each gene set’s constituent genes relative to the rest of the sample cohort.

### GSVA Heatmap visualization

GSVA enrichment scores were visualized as heatmaps using the Complex Heatmap package (v2.26.0) in R (v4.5.2). Pathways (rows) and samples (columns) were displayed in a fixed, biologically informed order with hierarchical clustering disabled on both axes. Columns were partitioned by disease status and, within each partition, ordered by additional sample-level covariates, all of which were rendered as categorical column annotations with fixed color palettes. Rows were partitioned by macro pathway category and annotated with both their macro- and micro-level classifications. Enrichment scores were mapped onto a divergent green–white–magenta color scale spanning the range of GSVA values. In addition to a heatmap of the full pathway set, per-category heatmaps were generated for a single time point, iterating over each macro category to produce individually scaled panels.

### Gene Set Enrichment Analysis (GSEA) FOUNDIN PD dataset and from PPMI/PDBP

Bulk RNA seq data from iPSC derived DA neurons from control and PD patients (n=98) were obtained from FOUNDIN-PD (PPMI, MJFF), expression data were obtained from salmon quants then processed for differential expression by Dseq2 and analyzed against a custom-curated gene set enrichment analysis for neurite and axon specific pathways obtained from MSigDB, Kegg and Hallmark pathway GSEA analysis performed using IDEP2 (South Dakota University) for blood transcriptomics processed Dseq2 data reanalyzed from the Parkinson’s Progression Markers Initiative (PPMI) and Parkinson’s Disease biomarkers discovery (PDBP) databanks published by Irmady et al ^31^. A false discovery rate (FDR) threshold of 0.25 was used to identify significantly enriched pathways.

### Proteomics data analysis

Protein intensity matrices were exported from Spectronaut (v20.6; library-free direct DIA; FDR 0.01 at precursor and protein levels). Known serum and media-derived contaminants were removed, and protein intensities were log₂-transformed. Proteins detected in at least 50% of samples in one experimental group were retained. Missing values were imputed using the MinProb method, followed by median centering. Protein identifiers were converted to gene symbols using clusterProfiler and org.Hs.eg.db. Differential protein expression between SMG and 1G conditions was assessed in 3D brain organoids and SH-SY5Y cells (n = 3/group) using the limma package with empirical Bayes moderation, and moderated t-statistics were used to rank proteins. For protein–protein interaction (PPI) analysis significance was defined as adjusted p < 0.05 and log₂FC ≥ 1.5-folds. PPI was performed using STRING v12 with a confidence threshold >0.40, incorporating experimental, curated database, co-expression, neighborhood, gene fusion, and co-occurrence evidence. Singleton nodes were removed. Network modules were identified using MCI clustering, and cluster-level functional enrichment was performed within STRING to identify overrepresented pathways ranked by enrichment significance^7,64^. Gene Set Enrichment Analysis (GSEA) was performed at Biomni^65^ using clusterProfiler (fgsea implementation) against Gene Ontology Biological Process, KEGG, and Reactome pathway databases. Pathways containing 15–500 genes were evaluated, and significance was determined using 10,000 permutations with Benjamini–Hochberg adjusted P < 0.05. Normalized enrichment scores (NES) were used to identify significantly enriched biological pathways.

### Inspiration4 PBMC transcriptomics analysis from SOMA browser

To assess spaceflight-associated transcriptional changes in proteostasis-related pathways, we interrogated peripheral blood mononuclear cell (PBMC) transcriptomic data from the Inspiration4 (I-4) mission using the Inspiration4 Multiome Data Explorer^32^. A curated list of ribosomal genes was first extracted from KEGG pathway annotations and used as a predefined gene set for downstream analysis. Gene-level expression of this ribosomal cohort was evaluated in astronaut PBMCs across pre-flight and post-flight time points. In parallel, we examined transcriptional regulation of the ubiquitin–proteasome system (UPS) by focusing on core components of proteostasis machinery, including ubiquitin genes, E3 ubiquitin ligases, and deubiquitinating enzymes (DUBs). Expression profiles of these UPS-associated genes were extracted from the same PBMC dataset and compared across flight phases to identify coordinated perturbations in protein synthesis and degradation pathways associated with spaceflight exposure.

### Statistical Analysis

All statistical analyses were performed in GraphPad Prism (v.9.5.1). Group comparisons used two-tailed unpaired Student’s *t*-test, with *P* < 0.05 considered statistically significant. Data are presented as mean +/- SEM unless otherwise indicated.

## Acknowledgment

Data used in the preparation of this article were obtained on [2024-05-30] from the Parkinson’s Progression Markers Initiative (PPMI) database (https://www.ppmi-info.org/access-data-specimens/download-data), RRID:SCR_006431. PPMI – a public-private partnership – is funded by the Michael J. Fox Foundation for Parkinson’s Research and funding partners, including AbbVie, Alamar Biosciences, Aligning Science Across Parkinson’s (ASAP), Arrowhead Pharma, Arvinas, AskBio, BIAL, BioArctic, Biohaven, BlueRock Therapeutics, Bristol Myers Squibb, Calico Labs, Capsida Biotherapeutics, Critical Path Institute, DaCapo Brainscience, Denali, Edmond J. Safra Foundation, Eli Lilly, Gain Therapeutics, GE Healthcare, Genentech, GSK, Insitro, Johnson & Johnson Innovative Medicine, Lundbeck, Merck, Neumora, Neuron23, Novartis, Olink, Regeneron, Roche, Sanofi, Tenvie, UCB, Vanqua Bio, Voyager Therapeutics, The Weston Family Foundation. For up-to-date information on the study, visit http://www.ppmi-info.org. We acknowledge Dr. Brad Morrison for kindly providing us with SH-SY5Y cell line. Schematic images were generated in Biorender.

## Author Contributions

Conceptualization: N.A.; Methodology and Investigation: N.A., SA., M.M., M.B., J.B., S.B., R.F.P., M.S., S.P.; Formal Analysis: NA; Writing—Original Draft: N.A., M.B., S.B.; Writing— Review & Editing: All authors; Visualization: N.A., S.A.; Funding Acquisition: N.A.; Supervision: N.A.

## Use of generative AI

During the preparation of this work, the authors used ChatGPT (OpenAI) for proofreading and language editing, including grammar, punctuation, and readability enhancement. After using this tool, the authors reviewed and edited the output as needed and take full responsibility for the content and accuracy of the final publication.

## Data Availability

All data generated are available for sharing upon request to NA. Patient data from PPMI is shared through PPMI data request process.

## Ethics Declaration

The study does not have any ethical declarations.

## Consent for Publication

All authors consented to the publication of this manuscript.

## References

1 Collaborators, G. B. D. P. s. D. Global, regional, and national burden of Parkinson’s disease, 1990-2016: a systematic analysis for the Global Burden of Disease Study 2016. Lancet Neurol 17, 939–953 (2018). 10.1016/S1474-4422(18)30295-3

2 Bloem, B. R., Okun, M. S. & Klein, C. Parkinson’s disease. Lancet 397, 2284–2303 (2021). 10.1016/S0140-6736(21)00218-X

3 Ascherio, A. & Schwarzschild, M. A. The epidemiology of Parkinson’s disease: risk factors and prevention. Lancet Neurol 15, 1257–1272 (2016). 10.1016/S1474-4422(16)30230-7

4 Lopez-Otin, C., Blasco, M. A., Partridge, L., Serrano, M. & Kroemer, G. The hallmarks of aging. Cell 153, 1194–1217 (2013). 10.1016/j.cell.2013.05.039

5 Lopez-Otin, C., Blasco, M. A., Partridge, L., Serrano, M. & Kroemer, G. Hallmarks of aging: An expanding universe. Cell 186, 243–278 (2023). 10.1016/j.cell.2022.11.001

6 Kalia, L. V. & Lang, A. E. Parkinson’s disease. Lancet 386, 896–912 (2015). 10.1016/S0140-6736(14)61393-3

7 Ali, N., Beheshti, A. & Hampikian, G. Space exploration and risk of Parkinson’s disease: a perspective review. NPJ Microgravity 11, 1 (2025). 10.1038/s41526-024-00457-6

8 Trovao, N. S. et al. in Fundamentals of Space Medicine and Clinical Technology (eds Ethan Waisberg, Joshua Ong, & Andrew G. Lee) 67-96 (Academic Press, 2026).

9 Chen, B. et al. The Impacts of Simulated Microgravity on Rat Brain Depended on Durations and Regions. Biomed Environ Sci 32, 496–507 (2019). 10.3967/bes2019.067

10 Hutchinson, A. M. et al. Advancing stem cell technologies for conservation of wildlife biodiversity. Development 151 (2024). 10.1242/dev.203116

11 Ellis, J. et al. Diversifying the reference iPSC line concept. Cell Stem Cell 32, 873–877 (2025). 10.1016/j.stem.2025.05.004

12 Jo, J. et al. Midbrain-like Organoids from Human Pluripotent Stem Cells Contain Functional Dopaminergic and Neuromelanin-Producing Neurons. Cell Stem Cell 19, 248–257 (2016). 10.1016/j.stem.2016.07.005

13 Smits, L. M. & Schwamborn, J. C. Midbrain Organoids: A New Tool to Investigate Parkinson’s Disease. Front Cell Dev Biol 8, 359 (2020). 10.3389/fcell.2020.00359

14 Marotta, D. et al. Effects of microgravity on human iPSC-derived neural organoids on the International Space Station. Stem Cells Transl Med 13, 1186–1197 (2024). 10.1093/stcltm/szae070

15 Pirjanian, N. A. et al. Establishing Neural Organoid Cultures for Investigating the Effects of Microgravity in Low-Earth Orbit (LEO). Methods Mol Biol 2951, 91–109 (2025). 10.1007/7651_2024_550

16 Bouten, C. V., Koekkoek, K. T., Verduin, M., Kodde, R. & Janssen, J. D. A triaxial accelerometer and portable data processing unit for the assessment of daily physical activity. IEEE Trans Biomed Eng 44, 136–147 (1997). 10.1109/10.554760

17 Thompson, M., Woods, K., Newberg, J., Oxford, J. T. & Uzer, G. Low-intensity vibration restores nuclear YAP levels and acute YAP nuclear shuttling in mesenchymal stem cells subjected to simulated microgravity. NPJ Microgravity 6, 35 (2020). 10.1038/s41526-020-00125-5

18 Herranz, R. et al. Ground-based facilities for simulation of microgravity: organism-specific recommendations for their use, and recommended terminology. Astrobiology 13, 1–17 (2013). 10.1089/ast.2012.0876

19 Klaus, D. M. Clinostats and bioreactors. Gravit Space Biol Bull 14, 55–64 (2001).

20 Hauslage, J., Cevik, V. & Hemmersbach, R. Pyrocystis noctiluca represents an excellent bioassay for shear forces induced in ground-based microgravity simulators (clinostat and random positioning machine). NPJ Microgravity 3, 12 (2017). 10.1038/s41526-017-0016-x

21 Dedolph, R. R. & Dipert, M. H. The physical basis of gravity stimulus nullification by clinostat rotation. Plant Physiol 47, 756–764 (1971). 10.1104/pp.47.6.756

22 Hammond, T. G. & Hammond, J. M. Optimized suspension culture: the rotating-wall vessel. Am J Physiol Renal Physiol 281, F12–25 (2001). 10.1152/ajprenal.2001.281.1.F12

23 Grimm, D. et al. Tissue Engineering Under Microgravity Conditions-Use of Stem Cells and Specialized Cells. Stem Cells Dev 27, 787–804 (2018). 10.1089/scd.2017.0242

24 Zigmond, M. J., Abercrombie, E. D., Berger, T. W., Grace, A. A. & Stricker, E. M. Compensations after lesions of central dopaminergic neurons: some clinical and basic implications. Trends Neurosci 13, 290–296 (1990). 10.1016/0166-2236(90)90112-n

25 Bezard, E., Gross, C. E. & Brotchie, J. M. Presymptomatic compensation in Parkinson’s disease is not dopamine-mediated. Trends Neurosci 26, 215–221 (2003). 10.1016/S0166-2236(03)00038-9

26 Ali, N. et al. 6-hydroxydopamine affects multiple pathways to induce cytotoxicity in differentiated LUHMES dopaminergic neurons. Neurochem Int 170, 105608 (2023). 10.1016/j.neuint.2023.105608

27 Chen, C. et al. Parkinson’s disease neurons exhibit alterations in mitochondrial quality control proteins. NPJ Parkinsons Dis 9, 120 (2023). 10.1038/s41531-023-00564-3

28 da Silveira, W. A. et al. Comprehensive Multi-omics Analysis Reveals Mitochondrial Stress as a Central Biological Hub for Spaceflight Impact. Cell 183, 1185–1201 e1120 (2020). 10.1016/j.cell.2020.11.002

29 Ali, N., Beheshti, A. & Hampikian, G. Unveiling Parkinson’s Disease-like Changes Triggered by Spaceflight. arXiv preprint arXiv:2408.15021 (2024).

30 Wakigawa, T. et al. Gravitational and mechanical forces shape mitochondrial translation. Nat Commun 17 (2026). 10.1038/s41467-026-74493-z

31 Irmady, K. et al. Blood transcriptomic signatures associated with molecular changes in the brain and clinical outcomes in Parkinson’s disease. Nat Commun 14, 3956 (2023). 10.1038/s41467-023-39652-6

32 Overbey, E. G. et al. The Space Omics and Medical Atlas (SOMA) and international astronaut biobank. Nature 632, 1145–1154 (2024). 10.1038/s41586-024-07639-y

33 Bressan, E. et al. The Foundational Data Initiative for Parkinson Disease: Enabling efficient translation from genetic maps to mechanism. Cell Genom 3, 100261 (2023). 10.1016/j.xgen.2023.100261

34 Yu, W., Sun, Y., Guo, S. & Lu, B. The PINK1/Parkin pathway regulates mitochondrial dynamics and function in mammalian hippocampal and dopaminergic neurons. Hum Mol Genet 20, 3227–3240 (2011). 10.1093/hmg/ddr235

35 Bonello, F. et al. LRRK2 impairs PINK1/Parkin-dependent mitophagy via its kinase activity: pathologic insights into Parkinson’s disease. Hum Mol Genet 28, 1645–1660 (2019). 10.1093/hmg/ddz004

36 Singh, F. & Ganley, I. G. Parkinson’s disease and mitophagy: an emerging role for LRRK2. Biochem Soc Trans 49, 551–562 (2021). 10.1042/BST20190236

37 Berg, D. et al. MDS research criteria for prodromal Parkinson’s disease. Mov Disord 30, 1600–1611 (2015). 10.1002/mds.26431

38 Harvey, E. J. & Ramji, D. P. Interferon-gamma and atherosclerosis: pro- or anti-atherogenic? Cardiovasc Res 67, 11–20 (2005). 10.1016/j.cardiores.2005.04.019

39 Zhu, Z. et al. Effects of dietary energy restriction on gene regulation in mammary epithelial cells. Cancer Res 67, 12018–12025 (2007). 10.1158/0008-5472.CAN-07-2834

40 Smits, P. et al. Lethal skeletal dysplasia in mice and humans lacking the golgin GMAP-210. N Engl J Med 362, 206–216 (2010). 10.1056/NEJMoa0900158

41 Meyer, G. F., Harrison, N. R. & Wuerger, S. M. The time course of auditory-visual processing of speech and body actions: evidence for the simultaneous activation of an extended neural network for semantic processing. Neuropsychologia 51, 1716–1725 (2013). 10.1016/j.neuropsychologia.2013.05.014

42 Surmeier, D. J., Obeso, J. A. & Halliday, G. M. Selective neuronal vulnerability in Parkinson disease. Nat Rev Neurosci 18, 101–113 (2017). 10.1038/nrn.2016.178

43 Ali, N. et al. A Study on Cybrid Model of Parkinson’s Disease from Indian Population. J Neurochem 118, 50–50 (2011).

44 Halton, T. L. et al. Low-carbohydrate-diet score and the risk of coronary heart disease in women. N Engl J Med 355, 1991–2002 (2006). 10.1056/NEJMoa055317

45 Enstero, M., Daniel, C., Wahlstedt, H., Major, F. & Ohman, M. Recognition and coupling of A-to-I edited sites are determined by the tertiary structure of the RNA. Nucleic Acids Res 37, 6916–6926 (2009). 10.1093/nar/gkp731

46 Wenceslau, C. F. & Rossoni, L. V. Rostafuroxin ameliorates endothelial dysfunction and oxidative stress in resistance arteries from deoxycorticosterone acetate-salt hypertensive rats: the role of Na+K+-ATPase/ cSRC pathway. J Hypertens 32, 542–554 (2014). 10.1097/HJH.0000000000000059

47 Zhao, Q. et al. Integrative multi-omics analysis reveals host-microbiome metabolic alterations and candidate biomarkers in Parkinson’s disease. BMC Microbiol (2026). 10.1186/s12866-026-05168-4

48 Swalley, S. E. Expanding therapeutic opportunities for neurodegenerative diseases: A perspective on the important role of phenotypic screening. Bioorg Med Chem 28, 115239 (2020). 10.1016/j.bmc.2019.115239

49 Doxakis, E. Insights into the multifaceted role of circular RNAs: implications for Parkinson’s disease pathogenesis and diagnosis. NPJ Parkinsons Dis 8, 7 (2022). 10.1038/s41531-021-00265-9

50 Cui, S. et al. Gene expression profiling analysis of locus coeruleus in idiopathic Parkinson’s disease by bioinformatics. Neurol Sci 36, 97–102 (2015). 10.1007/s10072-014-1889-z

51 Magadi, S. S. et al. Neuroinflammation and neurodegeneration trigger a specific splice form of ribosomal protein S24. Brain (2026). 10.1093/brain/awag166

52 Kim, J. W. et al. Defects in mRNA Translation in LRRK2-Mutant hiPSC-Derived Dopaminergic Neurons Lead to Dysregulated Calcium Homeostasis. Cell Stem Cell 27, 633–645 e637 (2020). 10.1016/j.stem.2020.08.002

53 Correddu, D. & Leung, I. K. H. Targeting mRNA translation in Parkinson’s disease. Drug Discov Today 24, 1295–1303 (2019). 10.1016/j.drudis.2019.04.003

54 Ashraf, D., Khan, M. R., Dawson, T. M. & Dawson, V. L. Protein Translation in the Pathogenesis of Parkinson’s Disease. Int J Mol Sci 25 (2024). 10.3390/ijms25042393

55 Moreno-Villanueva, M. et al. Transcriptomics analysis reveals potential mechanisms underlying mitochondrial dysfunction and T cell exhaustion in astronauts’ blood cells in space. Front Immunol 15, 1512578 (2024). 10.3389/fimmu.2024.1512578

56 Ao, X. et al. Longitudinal transcriptomic and epigenetic analysis of the blood in two astronauts. Sci Rep 15, 27175 (2025). 10.1038/s41598-025-13383-8

57 Clavel, M. A. et al. Association of B-Type Natriuretic Peptide With Survival in Patients With Degenerative Mitral Regurgitation. J Am Coll Cardiol 68, 1297–1307 (2016). 10.1016/j.jacc.2016.06.047

58 Balter, M. Human evolution. Ancient DNA from Siberia fingers a possible new human lineage. Science 327, 1566–1567 (2010). 10.1126/science.327.5973.1566-b

59 Gruss, L. T. & Schmitt, D. The evolution of the human pelvis: changing adaptations to bipedalism, obstetrics and thermoregulation. Philos Trans R Soc Lond B Biol Sci 370, 20140063 (2015). 10.1098/rstb.2014.0063

60 Zhang, Q. et al. Circulating mitochondrial DAMPs cause inflammatory responses to injury. Nature 464, 104–107 (2010). 10.1038/nature08780

61 West, A. P. et al. Mitochondrial DNA stress primes the antiviral innate immune response. Nature 520, 553–557 (2015). 10.1038/nature14156

62 Tansey, M. G. et al. Inflammation and immune dysfunction in Parkinson disease. Nat Rev Immunol 22, 657–673 (2022). 10.1038/s41577-022-00684-6

63 Sabate-Soler, S. et al. Released mitochondrial DNA and neurofilament light chain as Parkinson’s disease phenotypes in patient-specific midbrain assembloids. bioRxiv, 2025.2003.2028.645921 (2025). 10.1101/2025.03.28.645921

64 Ali, N. et al. 9S1R nullomer peptide induces mitochondrial pathology, metabolic suppression, and enhanced immune cell infiltration, in triple-negative breast cancer mouse model. Biomed Pharmacother 170, 115997 (2024). 10.1016/j.biopha.2023.115997

65 Huang, K. et al. Biomni: A General-Purpose Biomedical AI Agent. bioRxiv (2025). 10.1101/2025.05.30.656746

